# Coupled dynamics of predator group formation and prey populations

**DOI:** 10.64898/2025.12.26.696504

**Authors:** Talia Borofsky, Erol Akçay, Daniel Rubenstein, Gili Greenbaum, Simon Levin

**Affiliations:** Stanford University; University of Pennsylvania Department of Biology; Princeton University Department of Ecology and Evolutionary Biology; The Hebrew University of Jerusalem Department of Ecology and Evolutionary Biology; Princeton University High Meadows Environmental Institute

## Abstract

A diverse array of social carnivores, such as orcas and wolves, also hunt cooperatively, but it is not well understood how prey availability feeds back with the size of social carnivore groups. Cooperative hunting may expand predators’ access to food by allowing them to hunt prey they could not catch alone, but many predator groups are much larger than the group size that would maximize individual food intake. One promising explanation is that if individuals can freely leave and join groups, then predators behaving adaptively should continue to join a group until the fitness that they achieve in the group is the same as that of being solitary. Here, we investigate the stable group sizes that form in a population of predator groups when predators hunt two types of prey: big prey that is best hunted in groups and small prey that is best hunted alone. In conjunction, we track the dynamics of the distribution of predator group sizes when individuals make decisions about whether to join or leave groups based on their own fitness. We find that if individual predators are able to freely leave groups and pair up with other individuals, starting new groups, then group sizes at equilibrium are much smaller than expected, generally staying at or below the size that maximizes individual fitness. Furthermore, accounting for flexible group sizes leads to ecological conditions that are not predicted if predator group sizes are held constant; in some regions of the parameter space, predators partition resources by forming a bimodal distribution of group sizes, and as a result, create situations where an increase in one prey type helps the other prey population grow. This result has applications to management of carnivore-livestock conflict, implying that increasing populations of wild prey may protect livestock.

## 2 Introduction

Many predators, such as wolves, orcas, and sea otters, hunt cooperatively (C. Eisenberg, 2013; Estes et al., 1998; Valls et al., 2015). While cooperative hunting can increase the chances of capturing large prey and defending kills from scavengers (Grinsted et al., 2019; MacNulty et al., 2012; Vucetich et al., 2004), it also imposes costs—–primarily the need to share food. Understanding when and how predators choose to form hunting groups, and how predator group behavior alters the relationship between predator and prey populations, remains a central question in behavioral and theoretical ecology.

One force that may select for the formation of predator groups is the ability to capture large prey through cooperative hunting (e.g. Baird and Dill, 1996; Grinsted et al., 2019; Kao et al., 2020; Smith et al., 2012). Models of the evolution of pairs of animals hunting cooperatively (Borofsky et al., 2024; Mesterton-Gibbons and Hardy, 2021; Packer and Ruttan, 1988) confirm this theory if big prey is very large, very available, and if cooperative hunters have a mechanism to preferentially pair up with other cooperators. However, observed group sizes are often much larger than the group sizes that maximize hunting success or net intake (e.g. Fanshawe and Fitzgibbon, 1993; MacNulty et al., 2012; Packer et al., 1990), causing some to doubt whether catching big prey is a sufficient explanation for the formation of predator groups (Packer et al., 1990).

Increased hunting success could still explain observed group sizes that exceed what is optimal, particularly if predators decide to leave and join groups based on their own fitness. Pioneering models by Clark and Mangel (1984, 1986) and Sibly (1983) explained that predators making selfish decisions to join a group should continue to join a group even as it grows beyond the size that optimizes group members’ fitness, so long as the fitness that joiners would obtain in the group is greater than that which would be obtained alone. These early models implicitly assumed a sufficient supply of individuals able to join, no fission or fusion with other groups, and that groups cannot exclude individuals who want to join.

It is not known which predator group sizes will emerge if multiple groups are forming and splitting at the same time in response to prey availability. Previous theoretical work has examined distributions of animal group sizes (Durrett et al., 1999; Gueron and Levin, 1995; Lerch and Abbott, 2024). and a model by Lerch and Abbott (2024) even linked group dynamics to population dynamics. In these models, the group-membership decisions are not explicitly based on fitness consequences. It remains to be seen how fitness-based decisions, such as the decision to join a group if it increases one’s food intake, will alter the group size distributions predicted by these dynamic models, or whether the group sizes that emerge will exceed the group sizes that optimize fitness, as predicted by Clark and Mangel (1984, 1986) and Sibly (1983).

Another factor in determining group sizes is their impact on prey populations, which can feedback on the costs and benefits of being in different sized groups. Previous work has modeled how cooperative hunting influences the rate at which predators kill and thus how it structures population dynamics in ecological communities, holding group size fixed (Berec, 2010; Cosner et al., 1999; Fryxell et al., 2025, 2022, 2007; Teixeira Alves and Hilker, 2017). In particular, Fryxell et al. (2025, 2022, 2007) focused on lions, modeling per-capita prey consumption as a function of group size (i.e., by dividing the group hunting rate by group size), and found that cooperative hunting can stabilize predatorprey dynamics and promote coexistence between predators and their prey. Fryxell et al. assume that group size distributions are static and that larger groups do not have a higher hunting rate, but their results may not hold in situations where cooperative hunting improves hunting success or if group formation responds to prey availability. Borofsky et al. (2024) modeled predators hunting two prey types— big prey and small prey — where cooperative hunting provided a hunting advantage by allowing predators to capture big prey that was more profitable but with a a smaller population grown rate (following well known allometric relationships, see Damuth, 1987). They thus found that cooperative hunting could cause steep declines in prey, ultimately leading to the extinction of cooperative hunting and a population of only solitary hunters. However, predator groups and big prey populations may be more able to persist if predators can switch more easily between solitary and cooperative hunting, if predator populations change along with their prey, and if cooperatively hunting group sizes are larger than two.

Thus, collective decisions about predator group membership can create ecological dynamics that feed back into the fitness landscape of group membership itself, intertwining the dynamics of predator group formation and both predator and prey population sizes. We do not yet know how individual animals behaving adaptively—i.e., making choices to join or leave groups that they believe will increase their personal fitness—will affect prey populations, nor how the dynamics of different prey populations drive the formation of predator groups. We thus aim to investigate the ecological feedback of predator group formation and cooperative hunting by developing a model that bridges previously separate bodies of theory: models of the optimal and equilibrium group size without individual supply limitations, models of dynamic group size distributions, predator-prey models that account for cooperative hunting.

We begin by developing a dynamic model of group sizes. The process of group formation and dissolution follows the assumptions of Clark and Mangel (1984); Sibly (1983) that aimed to explain why predator groups are so large; individuals can freely leave or join groups that they encounter based on personal fitness. Our framework builds on the group fission-fusion model of Gueron and Levin (1995), extended to incorporate ecological feedback and adaptive individual decision-making.

We then extend this model to investigate predators hunting prey: first, we model predators hunting big prey that are best caught in groups; then, to examine how group size responds to prey choice, we introduce a small prey type that can be caught alone. Using this model, we ask whether the availability of big prey that are best caught in groups selects for predator group formation, and if so, whether it favors the formation of large groups—that is, groups that are larger than the group size that would maximize individual fitness. Then, we ask how dynamic group formation impacts the coexistence of the two types of prey by measuring apparent competition. Our results show that although predator group sizes tend to increase when large prey are more abundant or more rewarding, those group sizes are much smaller than we would expect. Notably, we sometimes observe bimodal group size distributions, with some predators remaining solitary while others form small groups. We also observe regions of parameter space where increasing big prey population sizes favor, rather than inhibit, population growth of small prey.

## 3 Model

We model a one-predator, two-prey system in continuous time, coupled with a model of the dynamics of the distribution of predator group sizes. We investigate this complex system by breaking it down into three stages. The first stage looks at dynamic group formation without population dynamics, where individuals decide whether to join or leave groups based on a fecundity function, *W*(*x*), which represents the benefits they would receive in a group of size *x*. The second stage examines cooperative hunting in a one-predator, one-prey system and then a one-predator, two-prey system, with all predators approximated as in groups of one size. The final stage merges the first two stages, defining fecundity as the benefit received from hunting in a group of size *x*.

### 3.1 Dynamic Group Size Distribution

We begin by modeling a population of constant size split into groups of size 1, 2, 3, …, *x*_*m*_, where *x*_*m*_ is the maximum group size and is a natural number. Although in biological reality, *x*_*m*_ could be arbitrarily large, in practice we assume that there is an upper limit on observed group sizes. The maximum group size can also be thought of as one more than the upper limit on the number of conspecifics with which an individual will tolerate living. Throughout the analysis, we set this maximum group size to be much larger than the observed most common and mean experienced group sizes at equilibrium, and thus groups at the maximum group size are so rare that they have a negligible effect on the shape of the group size distribution.

Fecundity, *W*(*x*), is the benefit that an individual receives in a group of size *x*, for *x* = 1, 2, 3, …, *x*_*m*_. Let *x*_0_ be the value of *x* that maximizes *W* (or the first value of *x* that maximizes *W*, for cases in which *W* has more than one maximum) and let *x*^*^ > 1 be the lowest group size at which it is no longer beneficial for a solitary individual to to join a group, i.e., *W*(*x*^*^) ≥ *W*(1) but *W*(*x*^*^ + 1) < *W*(1). For this section, we only assume that the fecundity function has an internal maximum; a specific functional form of *W* will be defined later.

Consider groups that form and dissolve from one individual at a time joining or leaving, where an individual is more likely to join a group if doing so increases its fecundity, and likewise, more likely to leave a group if doing so increases its fecundity. Clark and Mangel (1984, 1986); Sibly (1983) showed that if groups cannot reject individuals that want to join, then groups should grow past the optimal size of *x*_0_. In the model of Clark and Mangel (1986), the equilibrium group size should be the group size, *x*^*^, at which it would no longer be beneficial for individuals to join that group.

To model the decision-making process of predators as they consider whether to join or leave groups, we introduce the best response function *S*(*x, y*), which is the probability, upon encounter, that an individual in a group of size *y* will switch to a group of size *x*. The inputs of *S*(*x, y*) must satisfy 1 ≤ *y* ≤ *x*_*m*_ and 1 ≤ *x* ≤ *x*_*m*_ − 1 because groups are at most of size *x*_*m*_. A rational predator with perfect knowledge should switch from being in a group of size *y* to being in a group of size *x* with probability 1 if *W*(*x*) > *W*(*y*), with probability 1/2 if *W*(*x*) = *W*(*y*), and with probability 0 if *W*(*x*) < *W*(*y*). Since knowledge is not perfect, we assume that the best response function is a a smooth version of this step function, modeled as the logistic function,

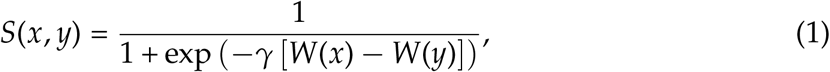

where *γ* is a positive scaling constant that determines the steepness of *S*(*x, y*), and can be thought of as the accuracy of the decision. Thus individuals have more certainty if the difference in fitness between being in a group of size *x* and *y* is larger; *S* is close to 0 if *W*(*y*) >> *W*(*x*), close to 1 if *W*(*x*) >> *W*(*y*), and somewhere between 0 and 1 if *W*(*x*) and *W*(*y*) are similar.

We can now explicitly formulate the rate of joining and leaving a particular group of size *x*. The time constant for group dynamics is *τ* (so 1/*τ* is the rate constant of group dynamics). If an individual encounters a group of size *x*, the rate at which it joins this group is 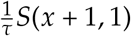 if *x* ≤ *x*_*m*_ − 1. The probability of joining the group is *S*(*x* + 1, 1) because if the individual joins, the group would be of size *x* + 1, and the fecundity that the individual would experience is *W*(*x* + 1). However, to add control over the rate at which pairs form, we weigh the rate at which a solitary individual pairs with a second solitary individual by the parameter *ϕ*, called the pairing parameter. Because we assume that groups do not control whether individuals leave or join, as in Clark and Mangel (1984, 1986); Sibly (1983), each individual in a group makes independent decisions of whether to leave the group. Thus the rate at which individuals leave a group of size *x* to be solitary is 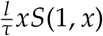 for *x* ≤ *x*_*m*_, where the parameter *l* is used to control the rate of leaving and called the leaving parameter.

Let *g*(*x, t*) be the number of individuals in a group of size *x* at time *t*, for *x* = 1, 2, …, *x*_*m*_, where we set a maximum group size because, for analysis, we need to limit the number of equations in the system. For ease of notation, we suppress dependence of *g* on *t* and let *g*_*x*_ = *g*(*x, t*). We assume predator groups move as units, and that each unit follows mass action laws, so the rate at which groups of size *x* encounter groups of size *y* is *g*(*x*)*g*(*y*), and thus the rate at which predators join groups of size *x* is 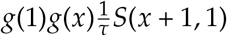. Note that the rate at which pairs form from solitary individuals that encounter each other must be multiplied by 1/2, to avoid double-counting of solitaries. The following system of *x*_*m*_ equations governs the continuous time dynamics of the distribution of group sizes:

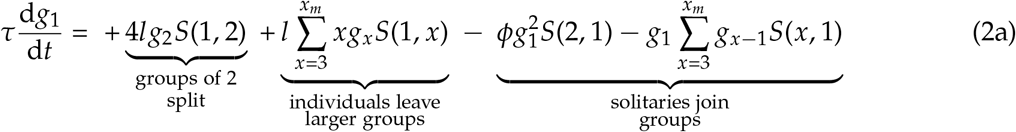

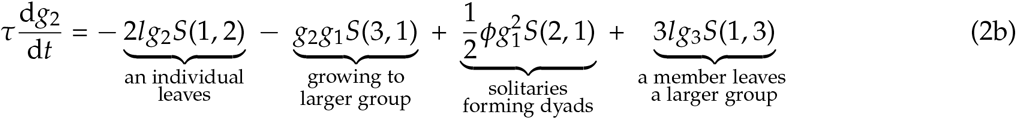

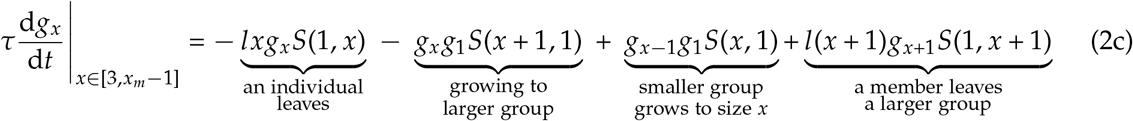

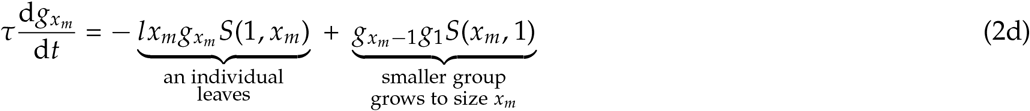

We use the mean experienced group size, 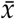, as a summary statistic of the group size distribution. If one chooses a random member of the predator population, 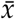 is the expected group size to which this individual belongs (also used in Niwa, 2003). In other words, 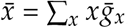, where 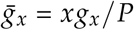 is the probability an individual is in a group of size *x* and *P* = ∑_*x*_ *xg*_*x*_. We do not use the mean group size, ∑_*x*_ *xg*_*x*_/∑_*x*_ *g*_*x*_, because it is biased toward small groups and does not tell us about the size of groups that individuals tend to be in; for example, if a population of 10 individuals is split into 3 singletons and one group of 7, the mean group size is 2.5, which makes it seem as if predators tend to be in small groups, when in fact most of them are in a large group.

In the main text, we set *l* = *ϕ* = 1, but examine the effect of modulating leaving and pairing in the appendix (Appendix Section A.1). We further examine robustness of the model in the appendix by introducing births and deaths, while maintaining constant population sizes (Appendix Section A.1.1). In this birth-and-death submodel, fecundity, *W*(*x*) is the birth rate per individual in a group of size *x*, and to keep the population size constant, we set the per capita death rate to *δ* = ∑_*x*_ *g*_*x*_*W*(*x*)/*P*.

### 3.2 Population Dynamics without Group Hunting

Here, we investigate the population dynamics of predators that cooperatively hunt prey in groups all of some constant size, *x*. We consider a predator population in a fixed area, with population size *P*. We will first consider the situation where it has one type of prey, big prey, with population size *M*_1_. We will then consider the situation where it can also hunt small prey, with population size *M*_2_ (we also refer to big prey and small prey as prey types 1 and 2, respectively). In this methods section, we present the model with both prey types and then outline how we explore the one-predator, one-prey submodel separately from the one-prey, two-prey model.

A predator group of size *x* kills prey of type *i* at rate *f*_*i*_(*x, M*_1_, *M*_2_), which is called the group functional response. The group then evenly shares the prey and each member converts prey caught to new (individual) predators at rate *b*_*i*_. Predators die at constant rate *δ*—note the redefinition of *δ* from the previous section. Each prey type exhibits logistic growth, with intrinsic growth rate *r*_*i*_ and carrying capacity *k*_*i*_. Thus, the population dynamics of predators and the two prey types are described by

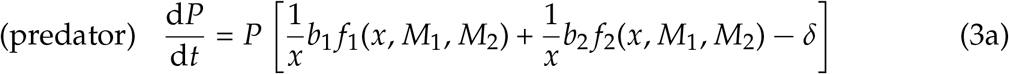

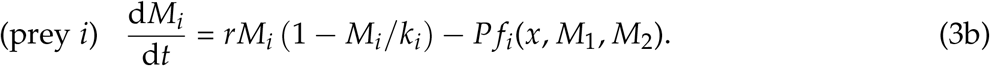

Assuming a predator hunting group (or lone predator) splits its time between collectively searching for prey and handling prey, we use a Type II functional response based on that of Holling (1959). For prey type *i*, the attack rate is decomposed into the encounter rate, *a*_*i*_, and the probability of capturing prey *i, α*_*i*_(*x*). As in Fryxell et al. (2007), the handling time is *h*_*i*_(*x*) = *h*_*ia*_ + *h*_*ib*_/*x*, where *h*_*ia*_ is the aspect of the handling time independent of group size and *h*_*ib*_ is the component of handling time that is split between the members of the group, assuming the split is even. The time *h*_*ib*_ can represent time spent consuming prey. Thus with both prey types present, the functional responses are,

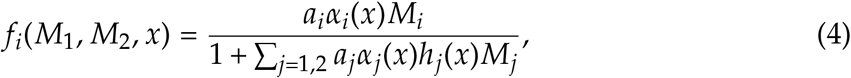

for prey types *i* = 1, 2.

The benefit of cooperation in the group is modeled as an increase in the probability of capturing large prey, so we assume that *α*_1_(*x*) is monotonically increasing in *x*. For small prey, we assume there is no benefit of cooperation on capture, so its capture probability *α*_2_(*x*) = *α*_2_, a constant. We further assume that capture probability of big prey accelerates with group size before leveling off, and thus use a sigmoidal function, namely,

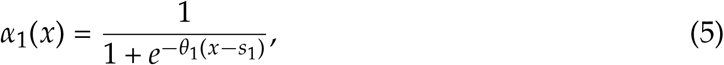

where the initial steepness of the curve is *θ*_1_ > 0, and *s*_1_ is the group size at which *α*_1_(*x*) = 1/2. To aid interpretation, we solve eq. 5 for *θ*_1_ in terms of *α*_1_(1), which yields 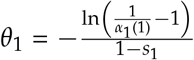. We assume a solitary predator is unlikely to catch big prey, so *α*_1_(1) is small.

Model parameters are listed in Table 1. To simplify parameters, we incorporate well-studied scaling laws relating population growth rates to body mass which have been applied to predator-prey models, as in Weitz and Levin (2006). As explained in Appendix Section B,

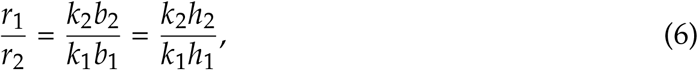

recalling that *r*_*i*_, *k*_*i*_, *b*_*i*_, and *h*_*i*_ are the intrinsic growth rate, carrying capacity, benefit to predators, and handling time of prey *i*, respectively. The relation between parameters in eq. 8 greatly increases tractability and focuses attention on the ratio between the benefit of big prey to small prey.

**Table 1.** List of parameters used in Project 1.

| Parameter | Definition |
| --- | --- |
| $r_i$ | predation-independent growth rate of prey type $i$ |
| $\delta$ | Death rate of predators, i.e. the probability a predator dies per unit ecological time; $\delta > 0$ |
| $k_i$ | carrying capacity of prey type $i$ |
| $b_i$ | Conversion of prey $i$ caught to predators; $b_1 > 0$ and $0 < b_2 < b_1$ |
| $\tau_x$ | The time constant for the dynamics of group size |
| $s_1$ | Critical group size for predators hunting prey type 1, i.e., the group size at which the capture probability of big prey is 1/2. $s_1 > 1$. |
| $h_{ia}$ | Group-size independent handling time of prey type $i$ by a hunting group of any size |
| $h_{ib}$ | Handling time of prey type $i$ by a single predator |
| $a_i$ | Attack rate on prey type $i$. |

#### 3.2.1 Non-dimensionalization

We partially non-dimensionalize the model, normalizing time and prey population sizes in order to reduce the number of parameters. The scaled prey sizes are *N*_1_ = *M*_1_/*k*_1_ for big prey and *N*_2_ = *M*_2_/*k*_2_ for small prey. The population time-scale constant is *ν* = *r*_1_ + *r*_2_ + *δ*, with non-dimensionalized time *T* = *tν*. The new parameters *T*_*g*_, *A*_*i*_, *H*_*ia*_, *H*_*ib*_, *β*_*i*_, *η*_*i*_ are defined in Table 2.

**Table 2.**
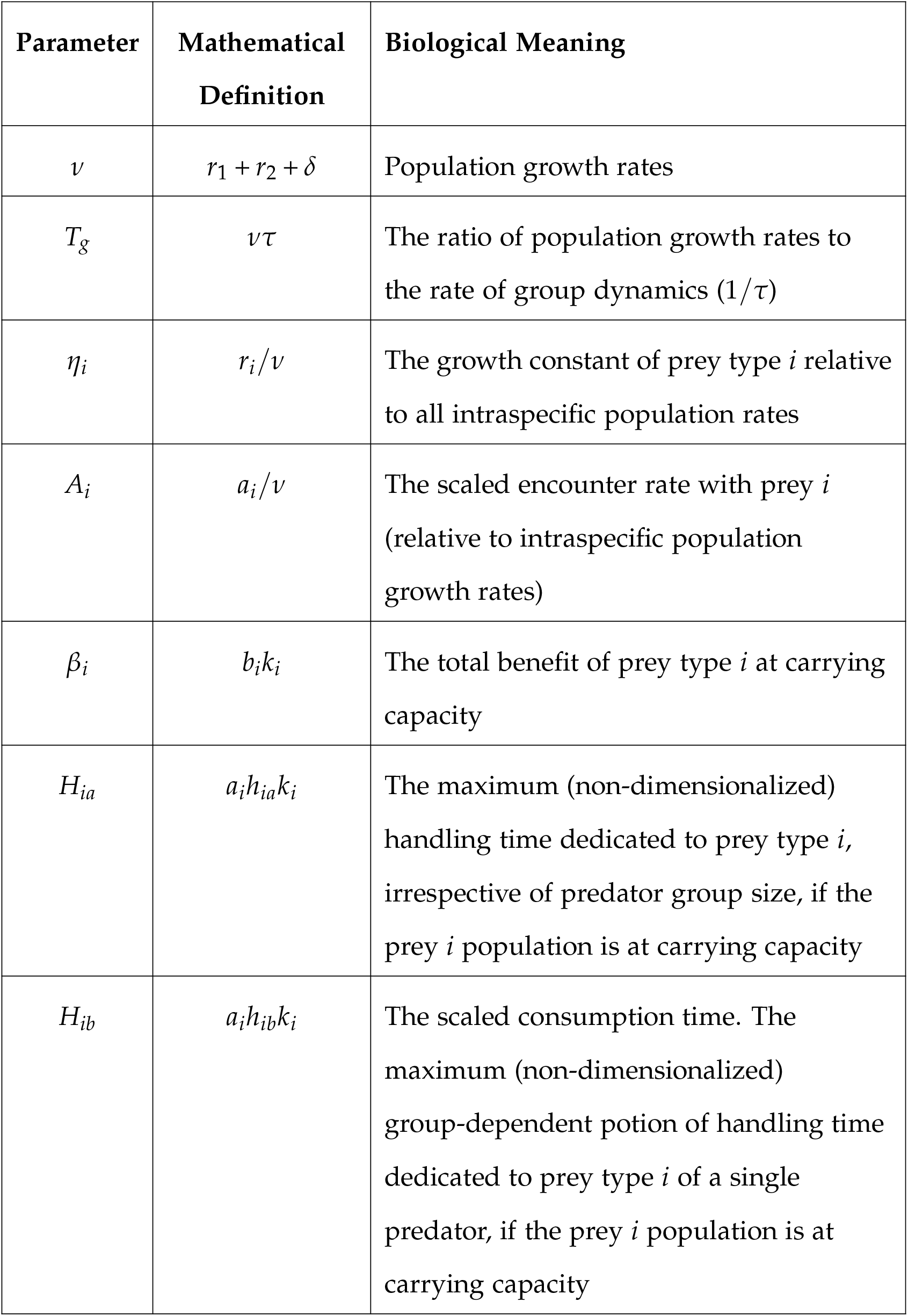
Parameters used for the non-dimensionalized model, their mathematical definitions in terms of the original parameters, and biological meanings.

Then the predator and prey population dynamics are

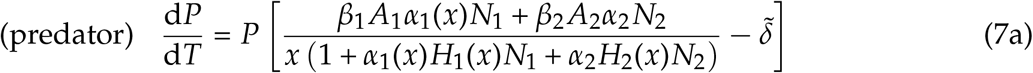

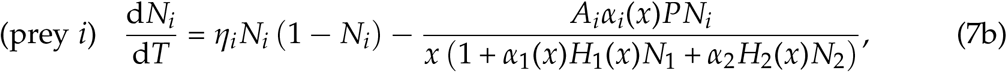

where 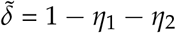.

Incorporating the scaling laws, as expressed in eq. 6,

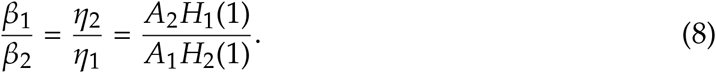

The term 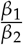, which we call the benefit ratio, is used frequently in subsequent analyses. From allometric scaling laws as explained in Appendix Sections B and because *β*_*i*_ = *b*_*i*_*k*_*i*_, for *m*_*i*_ the body mass of an individual of prey type *i* and *ϵ* a fraction such that 2/3 < *ϵ* < 1, we have *β*_1_/*β*_2_ = (*m*_1_/*m*_2_)^1−*ϵ*^, so the benefit ratio of the prey types grows as the mass ratio grows.

#### 3.2.2 Analyzing one-predator, one-prey and one-predator, two-prey separately

We first analyze this submodel without small prey and then with both types of prey present. The system governing the dynamics of predator and big prey, without small prey, is equivalent to setting *M*_2_ = 0 in eqs. 3a-b. The functional response to big prey is *f*_1_(*M*_1_, *x*) = *f*_1_(*M*_1_, 0, *x*) (see eq. 4). After non-dimensionalizing, the one-predator, one-prey system is 7a-b for *N*_2_ = 0. We specify small-prey associated parameters (e.g., *η*_2_) for this one prey population analysis in order to calculate big prey parameters (e.g., *η*_1_) using allometric scaling, and to maintain consistency in the non-dimensionalized parameters used across the two sub-models.

### 3.3 Merging Group Dynamics and Population Dynamics

This stage of the model links the previous stages, with predators split into groups of size *x* = 1, 2, …, *x*_*m*_ hunting one or two types of prey. As in the stage with only population dynamics, we analyze first the situation with only big prey available, and then with both big prey and small prey, but present in the methods the model with both prey types. Unlike Mesterton-Gibbons and Hardy (2021) and Borofsky et al. (2024), all predators are able to hunt alone or cooperatively in groups, so we focus on hunting group formation dynamics rather than the evolution of cooperative hunting. Recall that *g*_*x*_ = *g*(*x, t*) is the expected number of groups of size *x* at time *t*; therefore, the portion of predators that are part of a group of size *x* is 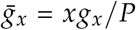 and *P* = Σ_*x*_*xg*_*x*_; hence *P* can also be thought of as the expected population size.

Under this formulation, the population dynamics of predators and both types of prey, altered from eqs. 3 to account for the group size distribution, can be described in part by the equations

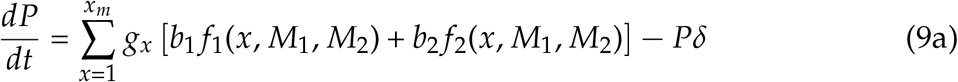

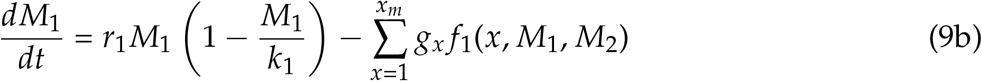

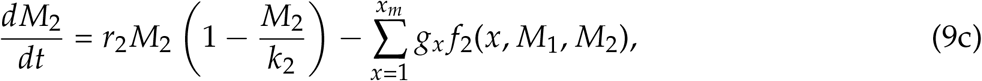

where the parameters are the same as in the previous section and are also summarized in Table 1. Eqs. 9a-c provide an incomplete description of the system, however, since they must be complemented by dynamic equations for the group sizes; those equations are altered from eqs. 2a - d as described below.

Joining and dispersing decisions, crucially, depend on their fecundities from hunting prey. Specifically, the fecundity of an individual in a group of size *x* is *W*(*x, M*_1_, *M*_2_), which is assumed to be a linear function of the yield from hunting. Fecundity is then defined as,

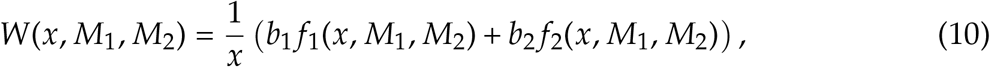

where we divide by *x* because we continue to assume the yield from hunting in a group is shared evenly. Note that *W*(*x, M*_1_, *M*_2_) is also the (per-capita) birth rate. For notational convenience, we henceforth suppress dependence on *M*_1_, *M*_2_. This new definition of the fecundity replaces *W* in the best response function, *S*(*x, y*) (eq. 1).

Recall that *τ* is the time constant for group dynamics. While time for the population dynamic processes in system eqs. 9 is on the order of years or seasons, time for group dynamics may be on the order of days, in which case *τ* << 1. The form of group joining and leaving rates are unaltered from eq. 2 (except *l* = *ϕ* = 1). The other two processes that change group size are birth and death. Recall that the rate at which individuals are born into a group of size *x* from birth is *xW*(*x*). Unlike in Section 1 and Appendix Section A.1.1, here the rate at which groups of size *x* shrink due to death is *xδ*, where *δ* is a constant.

Thus the master equations for the time evolution of *g*_*x*_, for *x* = 1, 2, …, *x*_*m*_, are

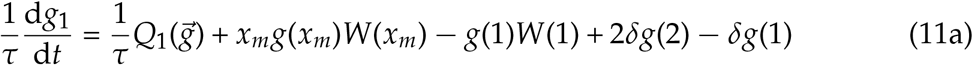

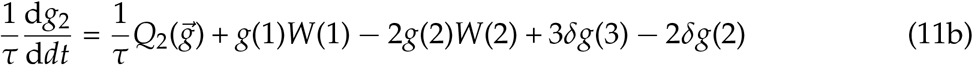

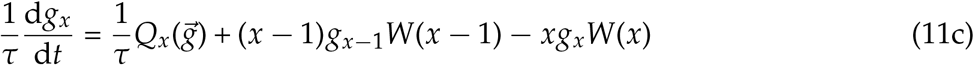

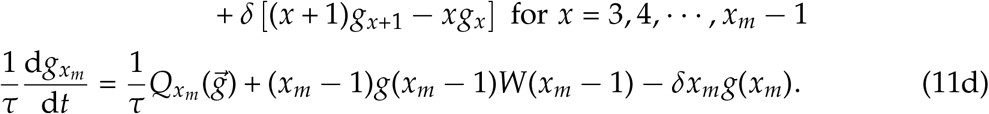

#### 3.3.1 Nondimensionalization and Rescaling

After non-dimensionalization as in Section 3.2.1, eqs. 9 become

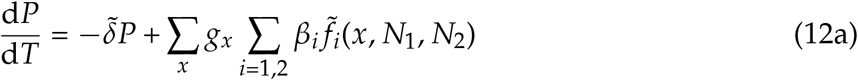

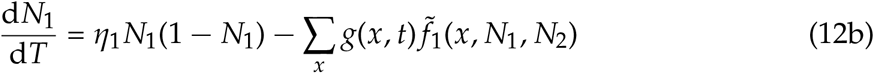

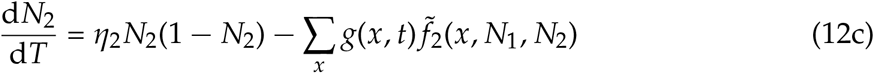

The rescaled functional response is 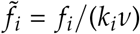 (see eq. C.23), so the rescaled fecundity is

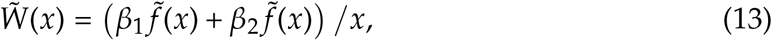

which is *W*(*x*)/*ν* (derived in Appendix Section C). Putting the best response function in terms of the non-dimensionalized variables and parameters, we replace *W* with 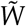 and *γ* with *d* = *γν* in eq. 1, (see eq. C.25 for the best response function, *S*(*x, y*), in terms of *d*).

In order to put the group dynamics in eqs. 11a-d in terms of *T* rather than *t*, we replace *W*(*x*) with 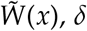 with 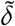, and *τ* with *T*_*g*_, so,

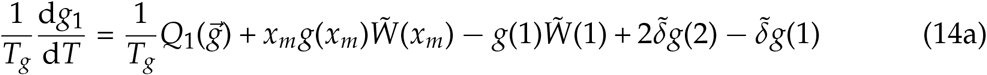

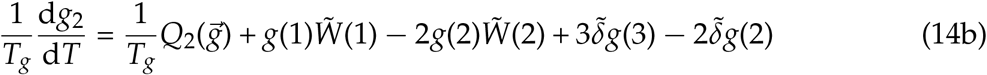

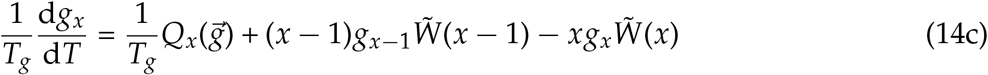

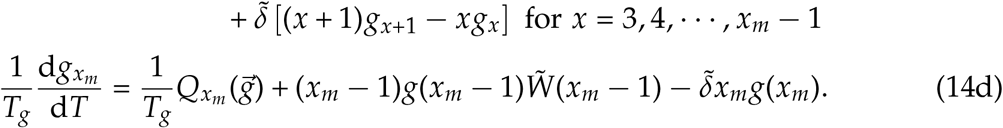

We do not non-dimensionalize predator population size or the group size distribution *g*_*x*_ because doing so does not help analysis and makes results more difficult to understand.

#### 3.3.2 One-predator, One-prey Submodel

As in the submodel without group dynamics, we first analyze the system with only one prey population, which is big prey, and then analyze the system with both prey populations. After non-dimensionalization, the one-predator, one-prey system is obtained by setting *N*_2_ = 0 in eqs. 12, paired with group dynamics in eqs. 14a-d. Fecundity, *W*(*x*) = *W*(*x, N*_1_) is defined in eq. 13 with *N*_2_ = 0. We continue to specify the parameters for small prey population dynamics in order to use allometric scaling in eq. 8 and the same non-dimensionalized parameters across submodels.

#### 3.3.3 Numerical Analysis

With one prey type and two prey types, the equilibria and eigenvalues cannot be solved analytically. The Jacobian for two prey present is in Appendix Section F.2, and the Jacobian for one prey present is obtained by eliminating the second column and row and setting *N*_2_ = 0. For a maximum group size of *x*_*m*_ = 10, we generated bifurcation diagrams (Figs F.15, F.16, F.17) using numerical continuation in Julia version 1.11.3 with the package BifurcationKit.jl (version 0.4.12; Veltz, 2020). The results of the numerical continuation are also used to plot the stable equilibria in Figs. 5, 6, and F.14. We generally set *α*_1_(1) = .05 (although see sensitivity analyses to *α*_1_(1) in Figs. F.18 and F.19). We also set the critical group size to *s* = 2 so that it is not too rare to have conditions that favor the formation of groups; a small increase in *s* to *s* = 2.45 can result in a requirement of very high abundances of very large prey for pairs to form (Fig. D.7), and larger groups cannot form without pairs forming first.

**Figure 1.**
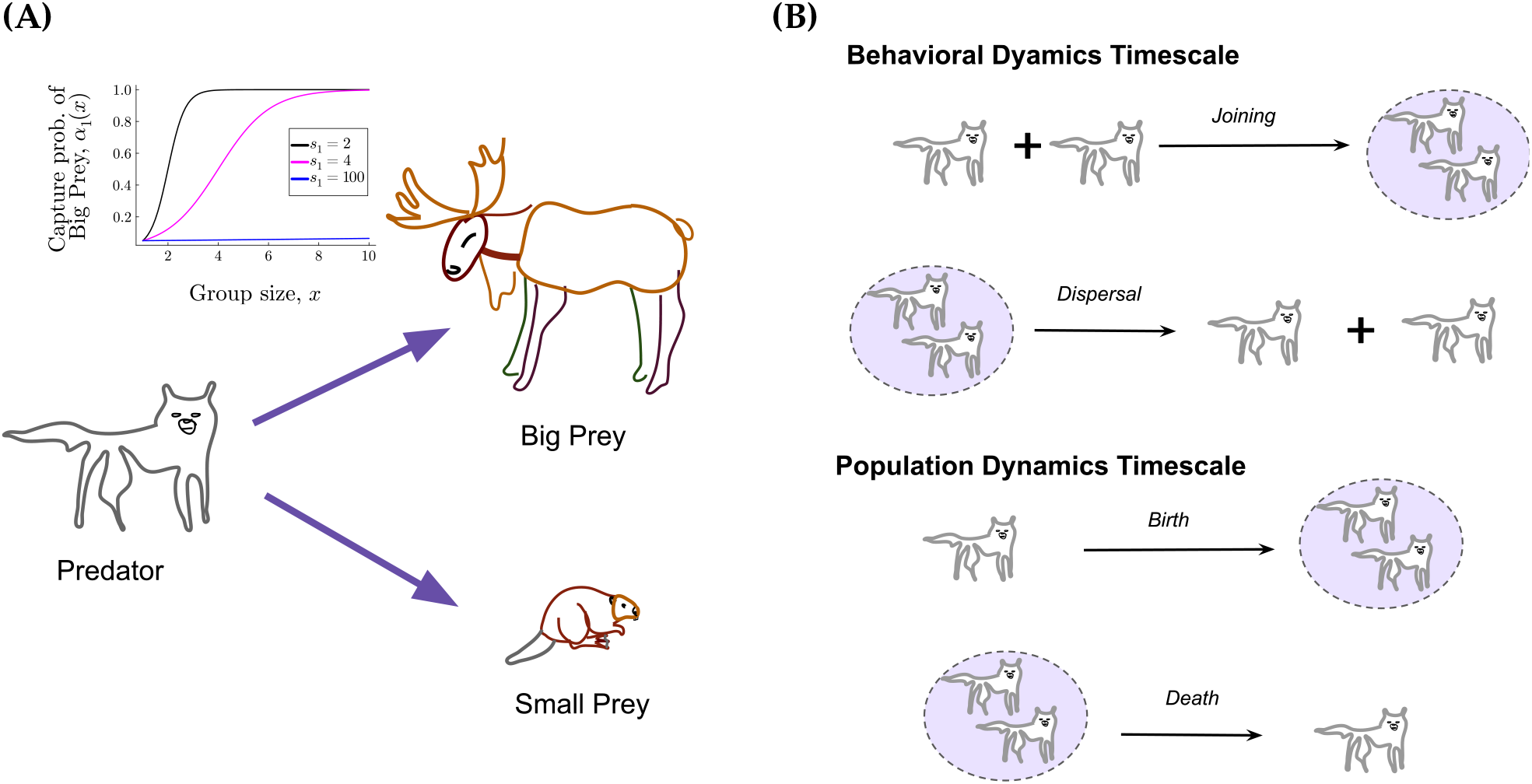
Diagram of the **A)** ecological, one-predator, two-prey portion of the model, and **B)** the group-dynamics. The capture probability of big prey versus group size is plotted panel **A)** for critical group size *s*_1_ = 2, 4, 100 and *α*_1_(1) = 0.05.

**Figure 2.**
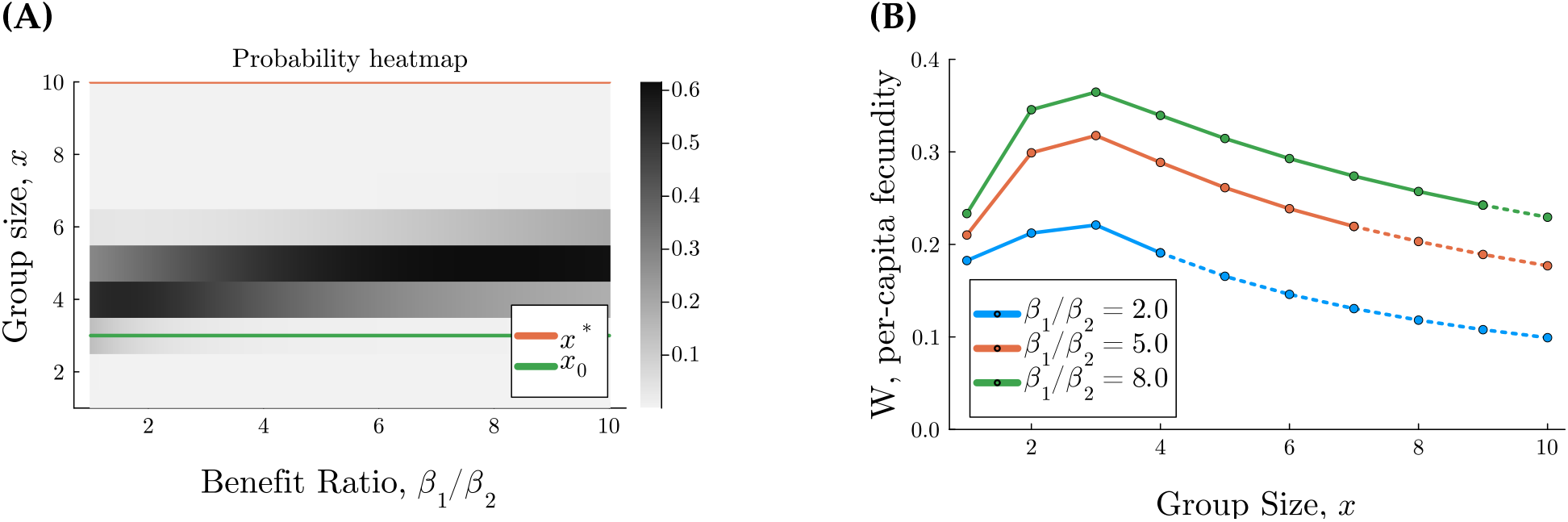
Group size distribution equilibrium when population sizes of predators and both prey are constant, with *P* = 10, *N*_1_ = 1, *N*_2_ = 0.5, and fecundity, 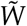, is based on functional responses to prey (eq. 13). Parameters are scaled by prey body mass such that *H*_1_(1)/*H*_2_(1) = *A*_1_*β*_1_/(*A*_2_*β*_2_). The parameters are *d* = 100, *H*_1*a*_ = *H*_2*a*_ = 0, *H*_2*b*_ = *β*_2_ = 1, *A*_1_ = *A*_2_ = 0.5, *s*_1_ = 2, *α*_1_(1) = 0.05, and *α*_2_ = 0.95 constant. Panel **A** shows the probability of experiencing a group of size *x* (indicated by the colorbar) at equilibrium versus *β*_1_/*β*_2_, with lines indicated the equilibrium group size predicted by Clark and Mangel (1986), *x*^*^ (orange), and the group size that maximizes fecundity, *x*_0_ (green). All equilibria are locally stable. Panel **C** shows the fecundity, *W*(*x*) versus group size, *x*, for different ratios *β*_1_/*β*_2_ (in color), with dotted lines connecting points where *W*(*x*) < *W*(1).

**Figure 3.**
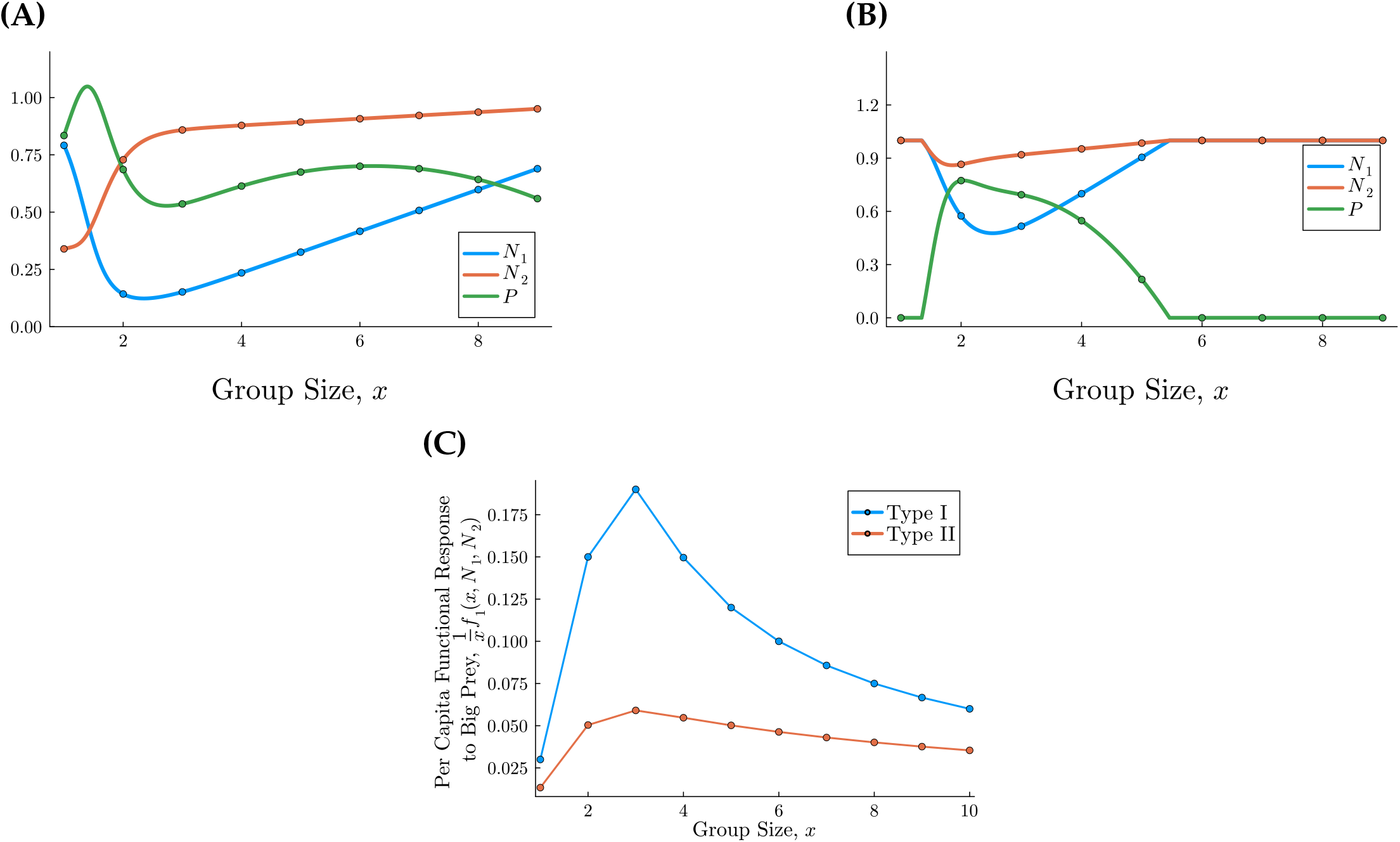
The effect of group size on the equilibrium predator, big prey, and small prey population sizes, without group dynamics (predators are in groups of the same size). Curves show stable equilibrium values of predator population size, i.e., *P* (green), and scaled big prey and small prey population sizes, i.e., *N*_1_, *N*_2_ (blue, red, respectively), for **A)** a type I functional response, i.e., *H*_1_(*x*) = *H*_2_(*x*) = 0, and **B)** a type II functional response, i.e., *H*_1_(*x*), *H*_2_(*x*) > 0. For both panels, the parameters use scaling laws (eq. 8), where the benefit ratio of prey is *β*_1_/*β*_2_ = 5.0, for *α*_1_(1) = 0.95, *α*_2_ = 0.05, *s*_1_ = 2, *η*_2_ = 0.6, and *β*_2_ = 1.0. For panel **B)**, *H*_1*a*_ = *H*_2*a*_ = 0 and *H*_2*b*_ = 1.0. Panel **C)** Shows the percapita functional response for the parameters in Panels **A, B**, which are blue and red, respectively. The continuous curves in panels **(A), (B)** come from numerical continuation, but the markers show that *x* is discrete.

**Figure 4.**
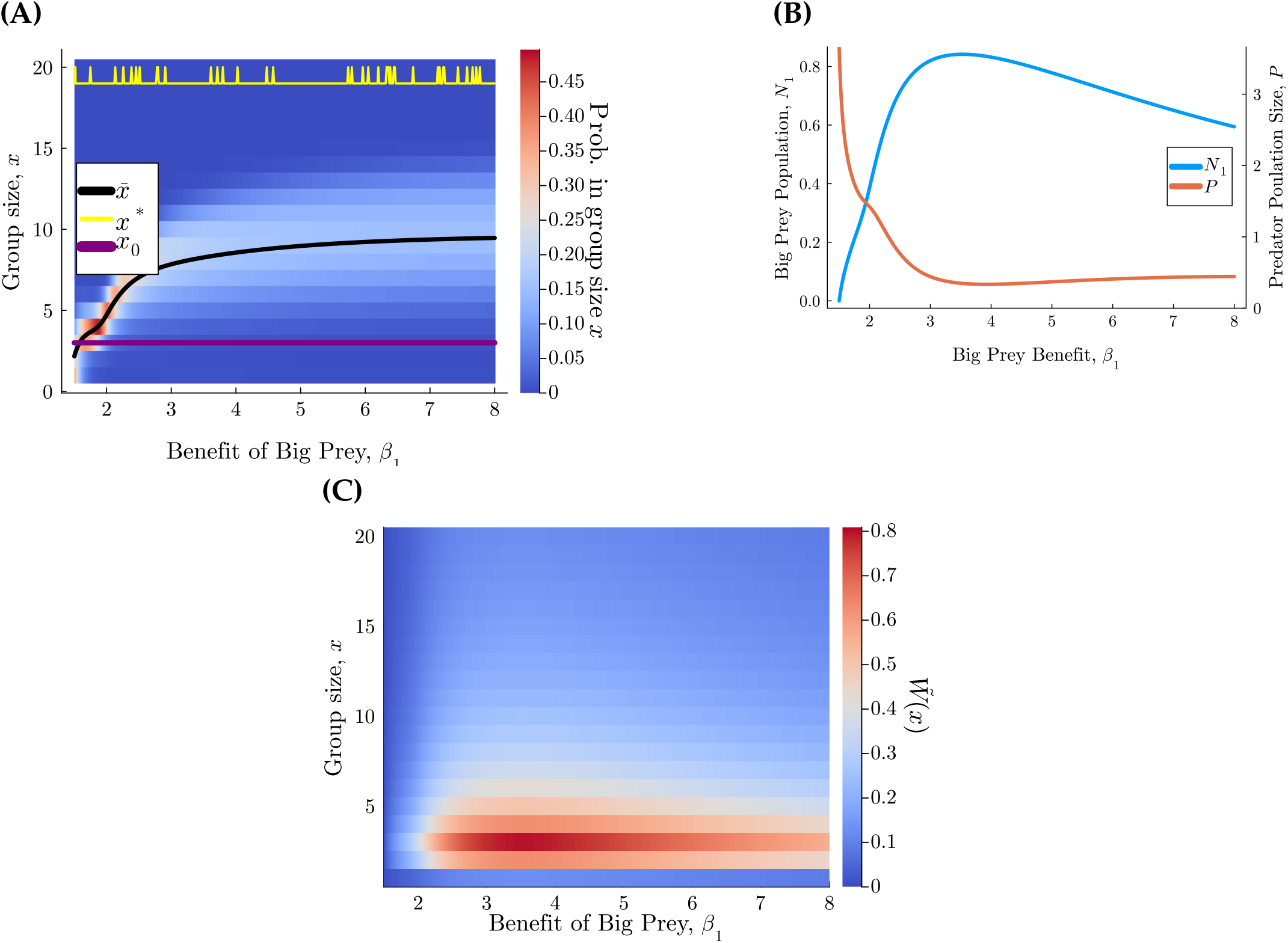
Increasing the benefit of big prey, *β*_1_ increases group size while decreasing predator population size, for the system with group dynamics and only one prey type. For varying *β*_1_ values (x-axis), panel **(A)** shows the probability of being in a group of size *x* (using a heatmap, as indicated by colorbar); panel **(B)** shows predator and big prey population sizes (*P* and *N*_1_, respectively); and panel **(C)** shows the fecundity at each group size, 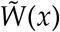, as indicated by a heatmap. The functional response is type I, i.e., *H*_1_(*x*) = 0. Parameters are allometrically scaled, where *x*_*m*_ = 20, *η*_2_ = 0.6, *β*_2_ = 1, *A*_1_ = 0.6, *A*_2_ = 0.5, *T*_*x*_ = 0.01, *d* = 100, *α*_1_(1) = 0.05, *s*_1_ = 2. In panel **(A)**, the purple line shows the group size that maximizes fecundity, the yellow line is the equilibrium group size predicted by Clark and Mangel (1986), and the black curve is the mean experienced group size. Spikes in the yellow line are artifacts of numerical instability because there are very small differences between *W*(19), *W*(20), and *W*(1).

**Figure 5.**
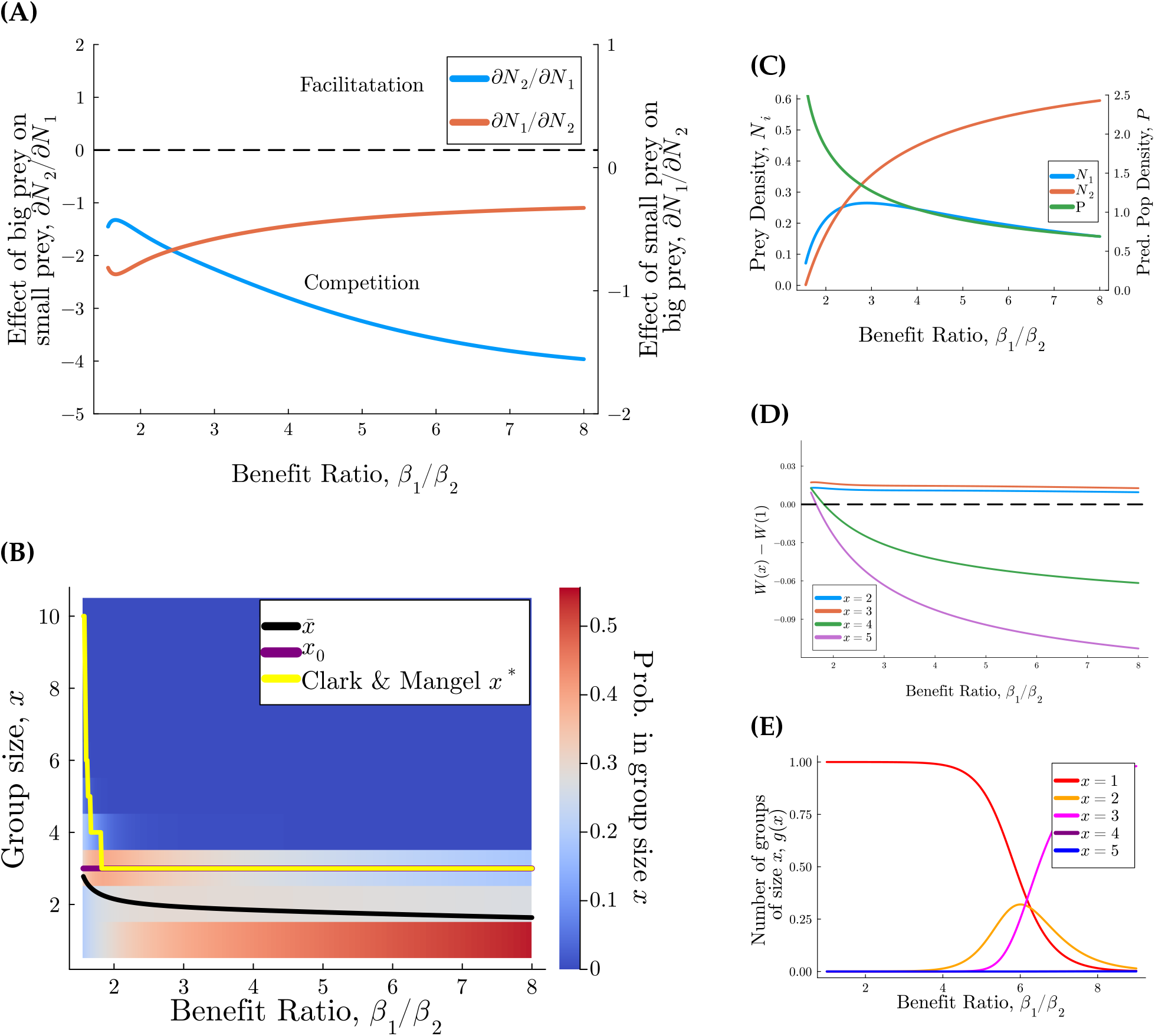
Characteristics of the stable coexistence equilibrium for group dynamics combined with population dynamics with a type I functional response, i.e., *H*_1_(*x*) = *H*_2_(*x*) = 0. (**A**) Apparent competition of prey 1 on prey 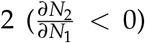 and of prey 2 on prey 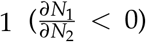. (**B**) The distribution of group sizes, with a heatmap indicating the proba-bility of experiencing a group of size *x, xg*(*x, t*)/*P*. The black curve is the mean experienced group size, 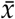, the yellow line is the equilibrium group size predicted by Clark and Mangel (1986), and the purple line is the group size that maximizes fecundity, *x*_0_. (**C**) Normalized population sizes of big and small prey, i.e., *N*_1_ and *N*_2_, respectively, and population size of predators, *P*. (**D**) Selection for predators to join or stay in larger group sizes, as represented by *W*(*x*) − *W*(1). (**E**) For comparison, the group size distribution at equilibrium if there are no population dynamics, for *N*_1_ = 0.2, *N*_2_ = 0.6, *P* = 3.0. The parameters are *α*_1_(1) = 0.05, *α*_2_(1) = 0.95, *s*_1_ = 2, *A*_1_ = 0.6, *A*_2_ = 0.5, *β*_2_ = 1.0, *η*_2_ = 0.6, *Tg* = 0.01, *x*_*max*_ = 10 and *d* = 100. The benefit of big prey and growth of big prey are found through the allometric scaling relationship in eq. 8.

**Figure 6.**
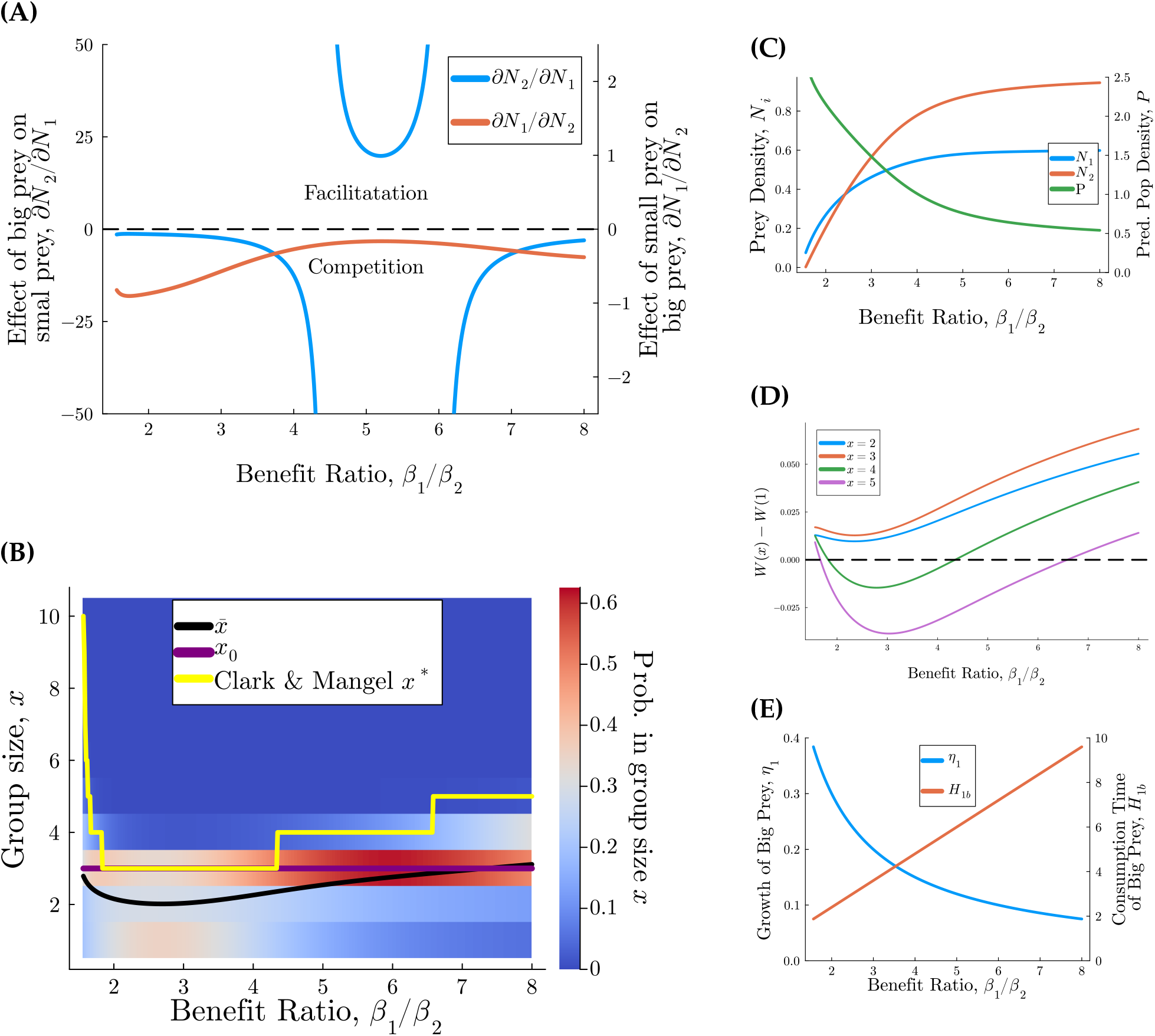
Characteristics of the stable coexistence equilibrium for group dynamics combined with population dynamics with a type II functional response, i.e., *H*_1_(*x*), *H*_2_(*x*) > 0. Parameters are allometrically scaled according to eq. 8. For each plot, the ratio of benefits of big prey to small prey, i.e., *β*_1_/*β*_2_, is varied. **(A)** The effect of prey 1 onprey 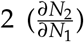, and vice versa 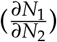, with regions of apparent facilitation and competi-tion. (**B**) The distribution of group sizes, with a heatmap indicating the probability of experiencing a group of size *x xg*(*x, t*)/*P*. The black curve is the mean experienced group size, 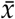, the yellow line is the equilibrium group size predicted by Clark and Mangel (1986), and the purple line is the group size that maximizes fecundity, *x*_0_. (**C**) Normalized population sizes of big and small prey, i.e., *N*_1_ and *N*_2_, respectively, and population size of predators, *P*. (**D**) Selection for predators to join or stay in larger group sizes, as represented by *W*(*x*) − *W*(1). **(E)** The relationship between consumption time, *H*_1*b*_, and scaled growth of big prey, *η*_1_ (eq. 8). The parameters are *s*_1_ = 2, *α*_1_(1) = 0.05, *α*_2_ = 0.95, *H*_1*a*_ = *H*_2*a*_ = 0, *H*_2*b*_ = 1.0, *A*_1_ = 0.6, *A*_2_ = 0.5, *T*_*g*_ = 0.01, and *d* = 100. The maximum group size is *x*_*m*_ = 10.

#### 3.3.4 Use of Generative A.I

Chat GPT (version 5.1) was used for editing of text for increased readability and for assistance with coding.

## 4 Results

We split analysis into first analyzing the time dynamics of the group size distribution, then population dynamics of predators and prey with predators split into groups of the same size, and finally the predator - prey model melded with the group dynamics equations (described in Sections 3.1, 3.2, and 3.3, respectively).

Throughout the results, we analyze the equilibrium group size distribution by comparing the mean experienced group size to the group size that maximizes fecundity, *x*_0_, and the predicted equilibrium group size predicted by Clark and Mangel (1986), called *x*^*^. We do this first with the group dynamics system where population size is restricted to be constant (eqs. A.2a-d and then with the group distribution equations combined with predator-prey population dynamics (i.e., eqs. 9a-c coupled with eqs. 14a-d). In addition, we compare the population sizes of predators and their prey, and the relationships between each population, both with and without dynamic group formation.

### 4.1 Group Dynamics with Constant Population Size

The group dynamics given in equation [2] have a unique equilibrium (Appendix Section A.2, Result 1). The behavior of the group size distribution at this equilibrium is shown in Figures 2 for *W* defined as in eq. [10] and in Appendix Figs. A.2 A.4, A.3, A.5, A.6 for alternative definitions of *W*. The mean experienced group size at this equilibrium is generally close to the group size, *x*_0_, that maximizes the fecundity of the group members, rather than *x*^*^, the group size at which it is no longer advantageous for a singleton to join.

We test the robustness of our results to incorporating births and deaths (while keeping population size constant) and suppression of leaving and pairing (Appendix Section A.1). Without suppression of leaving and pairing, i.e., *l* = *ϕ* = 1, adding birth and death seems to generally result in a noisier, dispersed final distribution. However, if leaving and pairing is suppressed to small rates (*l* = *ϕ* = 0.01), the combination with births and deaths leads to more groups of larger size, pushing 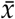 closer to *x*^*^ (Figs A.2, A.3, and A.4). When there are few opportunities to interact with singletons, births help push groups to larger sizes, particularly if offspring leave at a very slow rate.

### 4.2 Population Dynamics with Constant Group Size

Here we keep group size constant and assume that all predators are in groups of size *x* (see eqs. 7). The coexistence, predator extinction, big prey extinction, and small prey extinction equilibria can be found analytically and are in Appendix Section E.

#### 4.2.1 One-predator, One-prey

First, we examine the system with only one prey type, which is big prey. With a Type I functional response, increasing group size does not have a visible effect on predator or big prey population sizes (Fig. E.8), but with a Type II functional response, increasing the benefit of big prey, *β*_1_, generally narrows the range of group sizes for which predators can persist; for the parameters tested, as *β*_1_ increases, predators can only persist if they are in group sizes close to the group size that maximizes fecundity, *x*_0_ = 3 (Fig. E.9). Recall that *β*_1_ is assumed to be directly proportional to big prey body size. Due to allometric scaling of prey population growth and handling time (eq. 8), predators need to be in group sizes near *x*_0_ in order to persist if *β*_1_ is large because otherwise the increase in handling time and decrease in prey population growth from increasing big prey body size drives predators to extinction.

#### 4.2.2 One-predator, two-prey

We then analyze the system with both prey types: big prey and small prey. We examine the impact of increasing group size on the population sizes at equilibria, for 1 < *x* < 9, if the functional responses are Type I (i.e., there is no handling time, *H*_1_(*x*) = *H*_2_(*x*) = 0) (Fig. 3B). Predator population size is maximized for most of the population solitary, *x* ≈ 1.395, and is minimized at *x* ≈ 2.723 but does not change dramatically with group size as *x* increases past 3. The population size minimum occurs near the group size that maximizes the per capita functional response to big prey (*x* = 3; Fig. 3C), where the increased hunting efficiency drives big prey to low population levels. The total population size of prey generally increases with group size, and in particular, increasing group size benefits small prey.

If the functional response is Type II, there is a limited range of group sizes in which the predator population can persist (Fig. 3A), as in the system with one prey type. For the parameter values used in Fig. 3B, predators go extinct if the average group size is too small or too large, corresponding to group sizes at which the per-capita functional response to big prey is very small (Fig. 3C). Predators cannot persist if group sizes are too small because the capture probability of big prey is low (*α*_1_(1) = 0.05; but see sensitivity analyses of *α*_1_(1) in Appendix Figs. E.12, E.13) and the parameters used in Fig. 3B do not allow the predator population to survive if only subsisting on small prey. Predators also cannot persist of groups are much larger than is optimal because the population growth from killing big prey is too small; the benefit of big prey, *β*_1_, is diluted among group mates, and the increased kill rate from being in a group is not enough to compensate for both sharing and big prey’s high handling time. The predator population is maximized for *x* ≈ 2, where groups are large enough that they can sometimes catch big prey without imposing too much predation pressure on big prey, but at the same time, pairs do not incur too high of a cost of sharing when hunting small prey.

In addition to examining the effects of cooperative hunting and group size on population sizes, we also investigate how group formation influences the persistence and coexistence of prey species. We thus calculate apparent competition between the prey species; the effect of big prey population growth on small prey 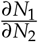, and small prey population growth on big prey, 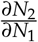, are calculated in Appendix Section E.4. We find that there is always apparent competition, rather than facilitation, between the two prey types (i.e.,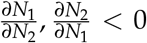) at the stable coexistence equilibrium if two conditions are met: first, the parameters follow the scaling relationships from Weitz and Levin (2006) as shown in eq. 8; and second, the group-independent portions of handling time for both types of prey are zero, i.e., *H*_1*a*_ = *H*_2*a*_ = 0 (see Result 7 in Appendix Section E.4).

### 4.3 Merging Group and Population Dynamics

#### 4.3.1 Analysis of equilibria, one prey type

If there is only one prey type present (big prey), then larger prey, i.e., increasing *β*_1_, favors larger groups (Fig. 4). The results are nearly identical for type I and type II functional responses (Fig. F.14). Mean experienced group size, 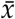, grows with *β*_1_ even though the group size that maximizes fecundity, *x*_0_, does not grow for the parameters tested. This is because the peak at *x*_0_ in the (scaled) fecundity function, 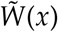, is less steep as *β*_1_ increases (Fig. 4c). In fact, for *β*_1_ large enough, 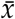 exceeds *x*_0_ by large margins, although it does not reach the group size predicted by **??**, *x*^*^.

#### 4.3.2 Analysis of predator joining decisions, two prey types

If given the opportunity to join a group, then big prey abundance, *N*_1_ and increased benefit relative to small prey, *β*_1_/*β*_2_ should favor individuals joining larger groups (i.e., *W*(*x*) > *W*(1) for larger *x*; shown in Appendix Section D). Increasing group size leads to a trade-off with hunting small prey, since it decreases the per capita functional response to small prey, 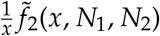 (Appendix Section D).

#### 4.3.3 Analysis of equilibria, two prey types

At the stable equilibria, increasing the body size of big prey (relative to small prey) does not necessarily lead to the formation of large hunting groups. If predators’ functional response is Type I (there is no handling time), then the mean experienced group size decreases with the ratio of big prey to small prey body benefit (Fig. 5B). Since the number of groups of size 3 and 4 increase as the ratio *β*_1_/*β*_2_ increases without population dynamics (Fig. 2), it seems that the low growth rate of big prey (see eq. 8), the low rate at which singletons encounter each other or other groups as the predator population size shrinks, and the constant rate of deaths prevent groups from growing.

If instead the functional response is Type II, mean experienced group size increases with the benefit ratio if the benefit ratio is high enough (Fig. 6B). This difference occurs because, as the benefit ratio *β*_1_/*β*_2_ increases, consumption time for big prey *H*_1*b*_ increases as well (Fig. 6E), but increasing group size decreases group consumption time, favoring larger groups. Furthermore, the Type II functional response limits predation pressure on big prey, as can be seen by the increased scaled population size of big prey, *N*_1_, with the Type II functional response (Fig. 6C) compared to *N*_1_ when predators have a Type I functional response (Fig. 5C).

We only observe bistability emerging if the rate of group dynamics is extremely fast, i.e., *T*_*g*_ < 10^−3^ (Fig. 6). For parameters such that 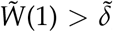, if *T*_*g*_ is very small, then we prove analytically that extinction of predators is stable (Appendix Section F.4.1).

For both Type I and II functional responses, the mean experienced group size is smaller than the equilibrium group size predicted by Clark and Mangel (1986), *x*^*^ (Figs. 5B, 6B). The mean experienced group size is also often smaller than the group size that maximizes fecundity, *x*_0_ (Figs. 5B, 6B).

It is surprising that predators are generally in groups smaller than even the group size that maximizes fecundity, which is not the case without population dynamics, for both types of fecundity functions and even with birth and death introduced (Figs. 2, A.2, A.3, A.4, A.5, A.6). This is also not the case in the system with only one prey type. However, in the regions where 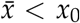, the optimal group size at the equilibria in both Figs. 5B, 6B has a very, very small fecundity advantage over being alone; in fact, *W*(*x*_0_) − *W*(1) < .02. Such a small difference means that the decision function, *S*(*x*_0_, 1), is not very close to one, and the rate of individuals leaving groups of size *x*_0_ is not rare. In fact, as *W*(*x*_0_) − *W*(1) increases, 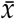 approaches *x*_0_ (see Fig. 6Bb,d). Furthermore, although at the actual equilibrium (for these parameters) groups of size *x*_0_ = 3 would maximize fecundity, because populations are dynamic, an increase in the number of groups of size *x*_0_ would lead to a further decrease in big prey—the less abundant of the two prey types — and an increase in small prey, which feeds back on the group size distribution by favoring solitary individuals.

Investigating apparent competition and the distribution of group sizes reveals that the system with dynamic group formation can behave quite differently from the system where all predators are in groups of the same constant size. The formulas for finding apparent competition of big prey onto small prey, 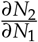, and small prey onto big prey, 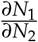, are in eqs. F.76, F.77, respectively. There exist regions in the parameter space (for the benefit ratio of big prey to small prey, *β*_1_/*β*_2_, at intermediate values) where big prey exerts apparent facilitation, rather than competition, on small prey(i.e., 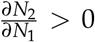; Fig 6a). The possibility of apparent competition here contrasts to the situation without group dynamics, where if *H*_1*a*_ = *H*_2*a*_ = 0 there is always bi-directional apparent competition between big prey and small prey (Result 7 in the Appendix). Additionally, the population cannot be well approximated as being split into groups around some mean experienced size 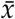; there exist regions where the group size distribution is bimodal, rather than unimodal (Fig. 6b).

## 5 Discussion

Our results support longstanding hypotheses that the availability of large prey, which are difficult to catch alone, favors the formation of hunting groups (e.g. Baird and Dill, 1996; Grinsted et al., 2019; Kao et al., 2020; Smith et al., 2012). We show that solitary predators are more likely to join larger groups, if given the opportunity, when the benefit ratio of big prey to small prey, *β*_1_/*β*_2_, and the availability of big prey, *N*_1_, are high. In both the systems with static and dynamic population sizes with a Type II functional response, the mean experienced group size increases as the benefit ratio rises, and solitary individuals become nearly absent from the population. In the full system coupling population and group dynamics, if there is only big prey present, then group sizes grow large, exceeding the group size that maximizes fecundity.

Nonetheless, with both prey types present, the resulting predator groups remain relatively small—generally around, or sometimes even slightly below, the group size that maximizes fecundity, or per-capita food intake. This contrasts with the results of Clark and Mangel (1986), who modeled the opportunity side of group formation, showing that oversized groups can emerge if individuals act selfishly—joining groups as long as it provides higher returns than hunting alone, even if group-level efficiency drops. However, our model—which also allows individuals to freely join or leave, but incorporates the supply side of group formation dynamics—produces groups that remain close to the size that maximizes individual intake.

The difference between our results and those of Clark and Mangel (1986) illustrates that the potential for formation of multiple groups lowers predicted group sizes. Clark and Mangel (1986) consider the case where singletons are deciding whether to join one growing group and there is no possibility of a second group. In our model, on the other hand, group sizes are eventually constrained by the supply of solitary individuals as mean group size increases. This constraint occurs because individuals are more likely to encounter each other or smaller groups and are thus rapidly siphoned away before they have the chance to join larger groups.

The dynamic nature of prey populations and thus the fitness landscape further restrains group sizes. With population dynamics merged with group dynamics and with both prey types present, the mean experienced group size at equilibrium is often smaller than even the group size that maximizes fecundity. Due to the fast population growth of small prey coupled with the slow regeneration of big prey, an increase in group sizes causes a rapid decline in the big prey population and an increase in the small prey population, favoring smaller predator groups.

Tweaks to the capture probability curve and group formation dynamics seem like they would promote the formation of larger groups but likely do not. For instance, if the capture probability curve were steeper (e.g., *s*_1_ > 2), larger groups might be more beneficial, but harder to form since intermediate-sized groups (like pairs) become rare. An alternative apparent solution to the problem of a dearth of singletons is to allow groups to merge and split apart, as in Gueron and Levin (1995), rather than only allowing one individual at a time to join a group. While altering our model to allow general mechanisms of group fission and fusion should be explored in future research, we do not expect it to lead to larger observed group sizes. Allowing groups to merge would require predators to compare the fitnesses between small and much larger groups, a scenario that typically disfavors merging due to the costs of sharing and saturating benefit of group size on hunting success. Similarly, allowing individuals to directly switch between groups should not drive up group sizes beyond those that optimize individual fitness.

Our result also contrasts with empirical observations of species like wild dogs and wolves, which often live and hunt in groups that exceed the size that maximizes per capita food intake (Fanshawe and Fitzgibbon, 1993; MacNulty et al., 2012; Thurber and Peterson, 1993)—although it is important to note that some have argued that accounting for energy constraints to hunting may yield higher optimal group sizes than previously calculated (Suter and Houston, 2020). Thus our results strongly suggest that for large groups to form, either there must be additional constraints on leaving groups and forming new groups, or the group size that optimizes fitness needs to increase.

Many group-living organisms have complex social structures that dictate the spatial structure of inter-group interactions (Creel and Creel, 2002; Jackson et al., 2017; Mech and Boitani, 2003; Mosser and Packer, 2009), which can suppress the rate at which individuals leave groups and start new ones. Territoriality and inter-group competition, in particular, can restrict the availability of territory in which individuals may settle and start new groups (Port et al., 2011; Zemel and Lubin, 1995). The limited supply of territories may constrain the number of smaller groups to join and lead to individuals staying longer in their natal group, causing existing groups to grow larger. Furthermore, territory supply could be closely related to prey availability, with existing wolf packs’ territories known to expand as prey populations drop (Dickie et al., 2022; Kittle et al., 2015; Prokopenko et al., 2024).

Additionally, large cooperative groups may evolve not solely due to hunting benefits, but instead synergy between multiple cooperative behaviors. Territorial defense, cooperative breeding, and hunting may co-evolve through reciprocal niche construction, where each behavior creates ecological conditions favoring the others (Van Dyken and Wade, 2012a,b). The group size that optimizes individual fitness may be much larger when considering these cooperative behaviors together rather than the one calculated when factoring in only the benefits of cooperative hunting. For example, Packer et al. (1990) concluded that access to large prey alone does not select for the observed size of lion hunting groups, and a more likely candidate is the need for defense in territorial encounters (Mosser and Packer, 2009). A phylogenetic review of Carnivora concluded that hunting of big prey was both significantly associated with the use of cooperative hunting and the presence of alloparental care (Smith et al., 2012). In fact, many well-known cooperative hunters—such as lions, wild dogs, wolves, and killer whales—also exhibit joint territorial defense or offspring care (Baird and Dill, 1996; Cassidy et al., 2015; Courchamp et al., 2002; Mosser and Packer, 2009).

Beyond the drivers of group formation, our model offers new insights into how social structure in predator populations influences ecosystem dynamics. Surprisingly, we found that group formation can benefit both prey types: prey populations generally increased with group size, while predator populations typically decreased. This contradicts models encoding cooperative hunting via Allee effects at the population level (Berec, 2010; Teixeira Alves and Hilker, 2017), but is consistent with models that modify the functional response using group size, not predator population size (Fryxell et al., 2022, 2007). It is interesting to note that this latter group of models by Fryxell et al. (2022, 2007) do not incorporate a benefit of cooperative hunting to hunting success. Thus, whether or not cooperative hunting boosts the capture probability of big prey, it decreases overall predation pressure on prey.

Dynamic group formation in predation also changes the relationships between prey populations. While constant group sizes always lead to bidirectional apparent competition between the two prey types if the group-independent portion of handling time, *H*_1*a*_ for big prey and *H*_2*a*_ for small prey, is negligibly small, the same is not true if group formation was dynamic; dynamic group formation creates regions of apparent facilitation. In these regions, increasing big prey population size reduces solitary predator numbers, which in turn alleviates predation pressure on small prey.

These insights could improve carnivore management strategies. As social carnivore populations, and particularly wolf populations, increase, so too does the incidence of livestock attacks (Chavez and Gese, 2005, 2006; Janeiro-Otero et al., 2020). Our model suggests that increasing abundance of alternative, wild prey that is hunted in groups may decrease the number of attacks on livestock. Furthermore, our model supports the assertion that by decreasing group sizes, lethal management of social carnivores that is meant to protect livestock might actually have the opposite effect (Imbert et al., 2016; Treves et al., 2024).

Ultimately, we show that integrating social dynamics into ecological models not only is feasible but also reveals critical mechanisms shaping species interactions. As recogni-tion of the ecological importance of social carnivores grows, such models will become essential tools for understanding and managing ecosystems.

## 6 Acknowledgements

We would like to thank Marcus W. Feldman for his comments on an earlier version of this manuscript. This study was supported in part in part by a gift by William H Miller III, the Agriculture Food Research Initiative (AFRI) Education and Workforce Development Fellowship, project award no. 2025-67012-45007, from the U.S. Department of Agriculture’s National Institute of Food and Agriculture, and and the Zuckerman STEM postdoc leadership program. Any opinions, findings, conclusions, or recommendations expressed in this publication are those of the authors and should not be construed to represent any official USDA or U.S. Government determination or policy.

## Notes

### Competing Interest Statement

The authors have declared no competing interest.

### Summary of Updates

We corrected the citation for BifurcationKit

