## Supplement for "Coupled dynamics of predator group formation and prey populations"

### A Group Dynamics Only

#### A.1 Robustness Analysis

We seek to understand why our results in the main text differ so greatly from the predictions of Clark and Mangel (1986), and test that the observed differences are robust to changes in the fecundity function.

We first specify the fecundity function, $W(x)$, using a gaussian-type function because it has parameters that easily control the location, width, and height of its maximum, which are called $x_0$, $\sigma$, and $a$, respectively (Fig. A.1). Then fecundity is

$$W(x) = ae^{-\frac{(x-x_0)^2}{2\sigma^2}}. \tag{A.1}$$

In subsequent sections, we will replace this fecundity function with one derived from the functional responses to prey.

The argument that group sizes should grow to the group size $x^*$ where fecundity is not improved by joining a group, i.e., $W(1) \geq W(x^*)$ (Clark and Mangel, 1986), assumes that there is no supply limitation on singletons available to join a growing group and that new groups do not form. However, if singletons follow mass action rules, then for an equal number of individuals that are solitary as are in groups of some size $x$, singletons should be more likely to interact with other singletons rather than with the groups. If singletons are also allowed to pair up, then the number of singletons available to join larger groups will decrease quickly. If $W(2) > W(1)$, then many singletons would quickly be siphoned away into pairs before they have a chance to join larger groups. The remaining singletons might then more quickly join the pairs to create groups of 3 than to join the group of size $x$ to create a group of size $x + 1$.

It thus seems that turning off the pairing of singletons by setting $\phi = 0$ in eqs. 2a-d should recover the prediction of Mangel and Clark. However, if we only turn off pairing, and start with one or more groups and some number $g_1$ of solitary individuals, then due to $S(1, x)$ being a smooth curve, individuals will occasionally, mistakenly leave the groups, and because no new groups are being made, the only equilibrium is for everyone to be in a group of size 1. We can prove this formally by taking the sum $\tau \sum_{x=2}^{x_m} \frac{dg(x)}{dt}$ for $\phi = 0$. This sum is equal to $-2lg_2S(1,2)$, so at equilibrium, $g_2 = 0$. By subsequently taking the sum $\tau \sum_{y=x}^{x_m} \frac{\mathrm{d}g(x)}{\mathrm{d}dt}$ for each $x > 2$, we find that at equilibrium each $g(x)$ is directly proportional to $g(x-1)$, and thus $g(x) = 0$ for $x \geq 2$. All individuals end up solitary. If we turn off leaving, this will stop $g_1^* = P$ from being an attractive state, but instead any $\vec{g}$ such that $g_1 = 0$ should be an equilibrium.

Thus, to try to recover the predictions of Mangel and Clark, we modulate both leaving of groups and the fusion of individuals into pairs by setting each of the parameters $l, \phi = 0.01$, a value chosen because it is very small. As shown in Figure A.2b, if the standard deviation of the fecundity curve is large, then very small leaving, pairing rates result in mean experienced group sizes that are closer to $x^*$ rather than $x_0$ (note $x_0 = 6$ in this figure). In contrast, with leaving and pairing unsuppressed, i.e, $l = \phi = 1$, the mean experienced group size is not only smaller than $x^*$ but also drops below $x_0$ if the standard deviation is high. However, for the standard deviation held constant at a low value of $\sigma = 1.0$, decreasing leaving and pairing has very little effect on $\bar{x}$, and $\bar{x}$ is generally closer to $x_0$ than to $x^*$ (Figures A.3, and A.4 panels a and b). The standard deviation, $\sigma$, of $W$ is important because if $\sigma$ is larger, then the fecundity of group sizes larger than $x_0$ are higher, increasing $S(x,1)$ for $x > x_0$. At the same time, increased $\sigma$ decreases the fecundity difference between being in a group and being alone, $W(x) - W(1)$, for intermediate group sizes, causing the probability individuals leave groups even if this decreases their fecundity to not be negligible.

##### *A.1.1 Births and Deaths with Constant Population Size*

We examine the effects of births and deaths on the group sizes. The number of births in a group is the fecundity function, $W(x)$, multiplied by the group size, i.e., $xW(x)$. To keep the population from growing out of control, we choose to keep it constant by setting the death rate per capita as the total birth rate divided by population size, i.e., $\delta = \sum_x g(x)W(x)/P$. Then the number of deaths in a group is $x\delta$. Assume that births cause a group size to be incremented by one unless the group is at the maximum group size, in which case each birth from a member of a group at the maximum size causes an increment in the number of solitary individuals.

Without births and deaths, let $\frac{dg_x}{dt} = Q_x(\vec{g})$, i.e., $Q_x(\vec{g})$ is the right side of eqs. 2 where $\vec{g} = (g_1, g_2, \cdots, g_{x_m})$. With births and deaths,

$$\frac{1}{\tau}\frac{dg_1}{dt} = \frac{1}{\tau}Q_1(\vec{g}) + x_m g_{x_m} W(x_m) - g_1 W(1) + 2\delta g_2 - \delta g_1 \tag{A.2a}$$

$$\frac{1}{\tau}\frac{dg_2}{ddt} = \frac{1}{\tau}Q_2(\vec{g}) + g_1 W(1) - 2g_2 W(2) + 3\delta g_3 - 2\delta g_2 \tag{A.2b}$$

$$\frac{1}{\tau}\frac{dg_x}{dt} = \frac{1}{\tau}Q_x(\vec{g}) + (x-1)g_{x-1}W(x-1) - xg_x W(x) \tag{A.2c}$$

$$+ \delta\left[(x+1)g_{x+1} - xg_x\right] \text{ for } x = 3, 4, \cdots, x_m - 1$$

$$\frac{1}{\tau}\frac{dg_{x_m}}{dt} = \frac{1}{\tau}Q_{x_m}(\vec{g}) + (x_m - 1)g_{x_m - 1}W(x_m - 1) - \delta x_m g_{x_m}. \tag{A.2d}$$

If pairing and leaving occur at the same rate as joining of groups, i.e., $\phi = l = 1$, then adding in birth and death appears to act more like noise rather than shifting the mean experienced group size. If instead leaving and pairing is suppressed, i.e., $\phi = l = 0.01$, then the addition of births and deaths results in mean experienced group sizes that are larger, and sometimes close to $x^*$ (Figs. A.2, A.3, A.4). Larger groups emerge because if pairing is rare, then births can help increase group size before too many solitary individuals are siphoned into smaller groups. Although deaths decrease group size, they can also help generate solitary individuals, particularly if leaving is suppressed.

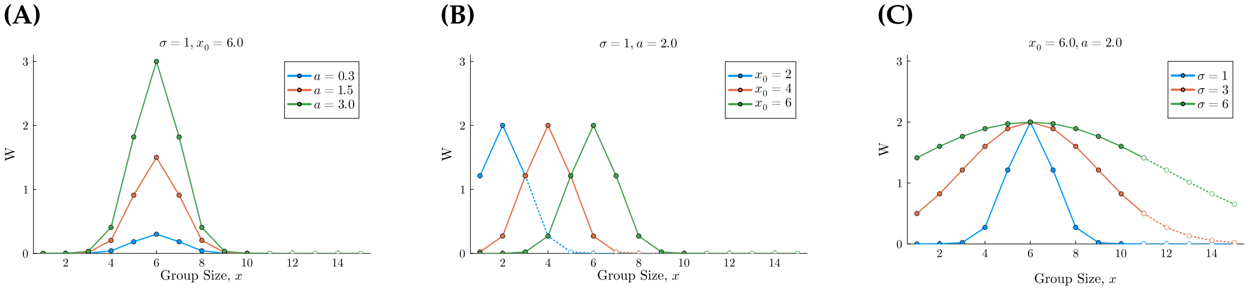


**Figure A.1:** Gaussian function for fecundity, $W(x)$ (y-axis) versus group size, $x$ (x-axis) with fecundity maximized at $x_0$, with standard deviation $\sigma$ and height $a$. Panels **(A), (B), (C)** show $W(x)$ for different values of $a, x_0$, and $\sigma$, respectively.

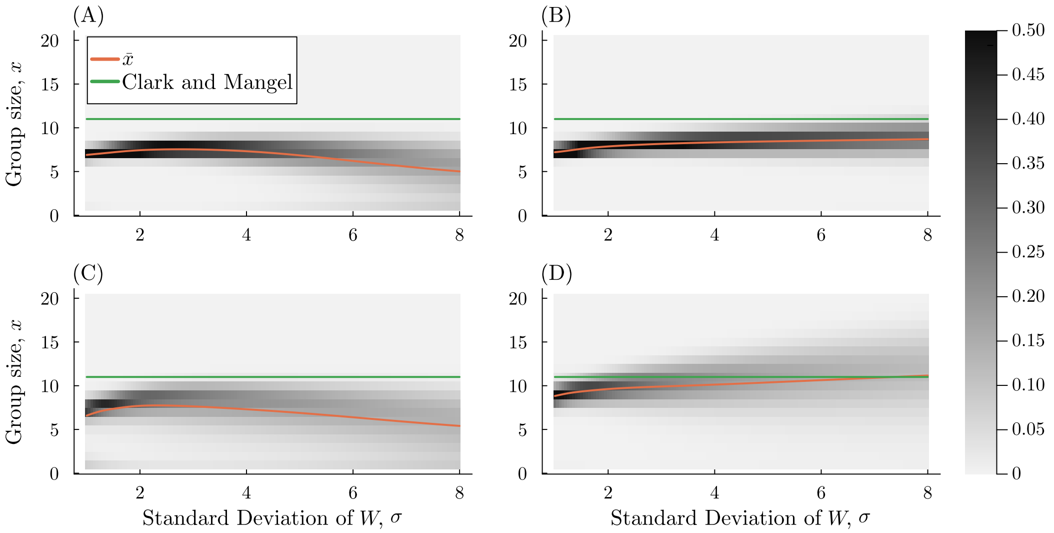


**Figure A.2:** Heatmap of the probability of being in a group of each size at equilibrium (indicated by the colorbar; calculated as $xg_x/P$ for $P$ the population size), as the standard deviation of the fecundity curve $\sigma$ varies (x-axis). The orange curve indicates the mean experienced group size, and the green line is $x^*$, the largest group size for which $W(x^*) \geq W(1)$, which is the equilibrium group size predicted by Clark and Mangel (1986). The top row (panels (A) and (B)) shows group dynamics without birth and death, whereas the bottom row (panels (C) and (D)) shows group dynamics with birth and death, for $\tau = 0.1$, where population size remains constant. For the left column (panels (A) and (C)), leaving and pairing of singletons occurs at the same rate as other group joining decisions (i.e. $l = \phi = 1$) and for the right column (panels (B) and (D)) leaving and pairing of singletons occurs very slowly, i.e. $l = \phi = 0.01$. For all panels, the maximum group size is $x_m = 20$, the steepness of the decision curve is $\gamma = 5$, fecundity is maximized at $x_0 = 6$, and the maximum fecundity is $a = 2.0$. Whereas we can solve for the equilibrium in the top row, for the bottom row, where it cannot be solved analytically, the starting point of all panels is $g_1 = 3.0, g_3 = 1.0, g_5 = 1.0$, and $g_x = 0$ otherwise, and the final time of the simulation is $50,000$. Starting at other initial points yields visually identical results.

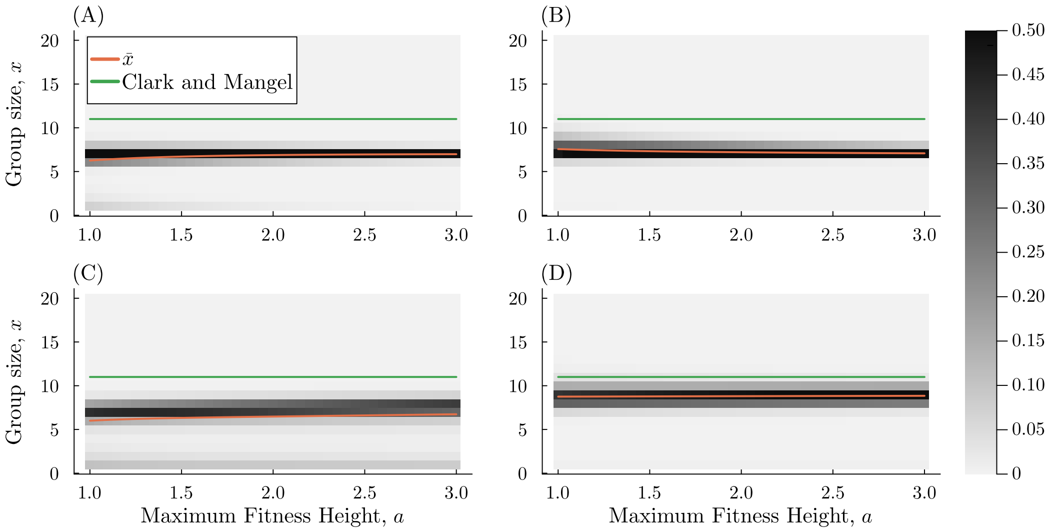


**Figure A.3:** Heatmap of the probability of being in a group of size $x$ at equilibrium (indicated by the colorbar), as the maximum fitness, $a$, varies (x-axis). The orange curve indicates the mean experienced group size, and the green line is $x^*$, the largest group size for which $W(x^*) \geq W(1)$, which is the equilibrium group size predicted by Clark and Mangel (1986). The top row (panels (A) and (B)) shows group dynamics without birth and death, whereas the bottom row (panels (C) and (D)) shows group dynamics with birth and death, for group dynamics time constant $\tau = 0.1$, where population size remains constant. For the left column (panels (A) and (C)), leaving and pairing of singletons occurs at the same rate as other group joining decisions (i.e. $l = \phi = 1$) and for the right column (panels (B) and (D)) leaving and pairing of singletons occurs very slowly, i.e. $l = \phi = 0.01$. For all panels, the maximum group size is $x_m = 20$, the steepness of the decision curve is $\gamma = 5$, fecundity is maximized at $x_0 = 6$, and the standard deviation if $\sigma = 1.0$. We solve for the equilibrium in the top row, but for the bottom row, we cannot solve for it analytically, so instead we iterate from a starting point. The starting point of all panels in the bottom row is $g_1 = 3.0, g_3 = w.0, g_5 = 1.0$, and $g_x = 0$ otherwise, and the final time of the simulation was $50,000$. Starting at other initial points yields visually identical results.

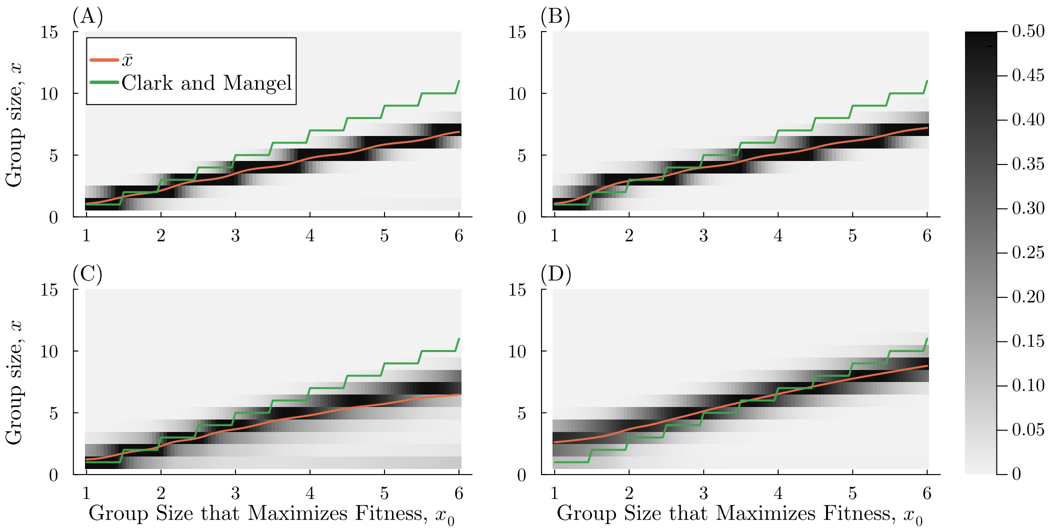


**Figure A.4:** Heatmap of the probability of being in a group of size $x$ at equilibrium (indicated by the colorbar), as the group size that maximizes fecundity, $x_0$, varies (x-axis). The orange curve indicates the mean experienced group size, and the green line is $x^*$, the largest group size for which $W(x^*) \geq W(1)$, which is the equilibrium group size predicted by Clark and Mangel (1986). The top row (panels (A) and (B)) shows group dynamics without birth and death, whereas the bottom row (panels (C) and (D)) shows group dynamics with birth and death, for group dynamics time constant $\tau = 0.1$, where population size remains constant. For the left column (panels (A) and (C)), leaving and pairing of singletons occurs at the same rate as other group joining decisions (i.e. $l = \phi = 1$) and for the right column (panels (B) and (D)) leaving and pairing of singletons occurs very slowly, i.e. $l = \phi = 0.01$. For all panels, the maximum group size is $x_m = 20$, the steepness of the decision curve is $\gamma = 5$, the maximum fecundity is $a = 2.0$, and the standard deviation is $\sigma = 1.0$. We solve for the equilibrium in the top row, but for the bottom row, we cannot solve for it analytically, so instead we iterate from a starting point. The starting point of all panels in the bottom row is $g_1 = 3.0, g_3 = w.0, g_5 = 1.0$, and $g_x = 0$ otherwise, and the final time of the simulation is $50,000$. Starting at other initial points yields visually identical results.

##### *A.1.2 Plots with Functional-Response Based Fecundity*

To further understand how the shape of the fecundity function affects the group size equilibria, we use a definition of fecundity based on the functional responses to prey, as in eq. 13, but simplified by setting $\beta_1 = 1, H_1(x) = 0, N_1 = 1$, and $N_2 = 0$. This simplified fecundity is then,

$$\tilde{W}(x) = \frac{1}{x} A_1 \alpha_1(x). \tag{A.3}$$

Using this definition of fecundity, groups can grow larger than the group size which maximizes per capita fitness, $x_0$, but they generally do not grow up until $W(x) \leq W(1)$, which occurs at group size $x^*$ (FIgs A.5, A.6).

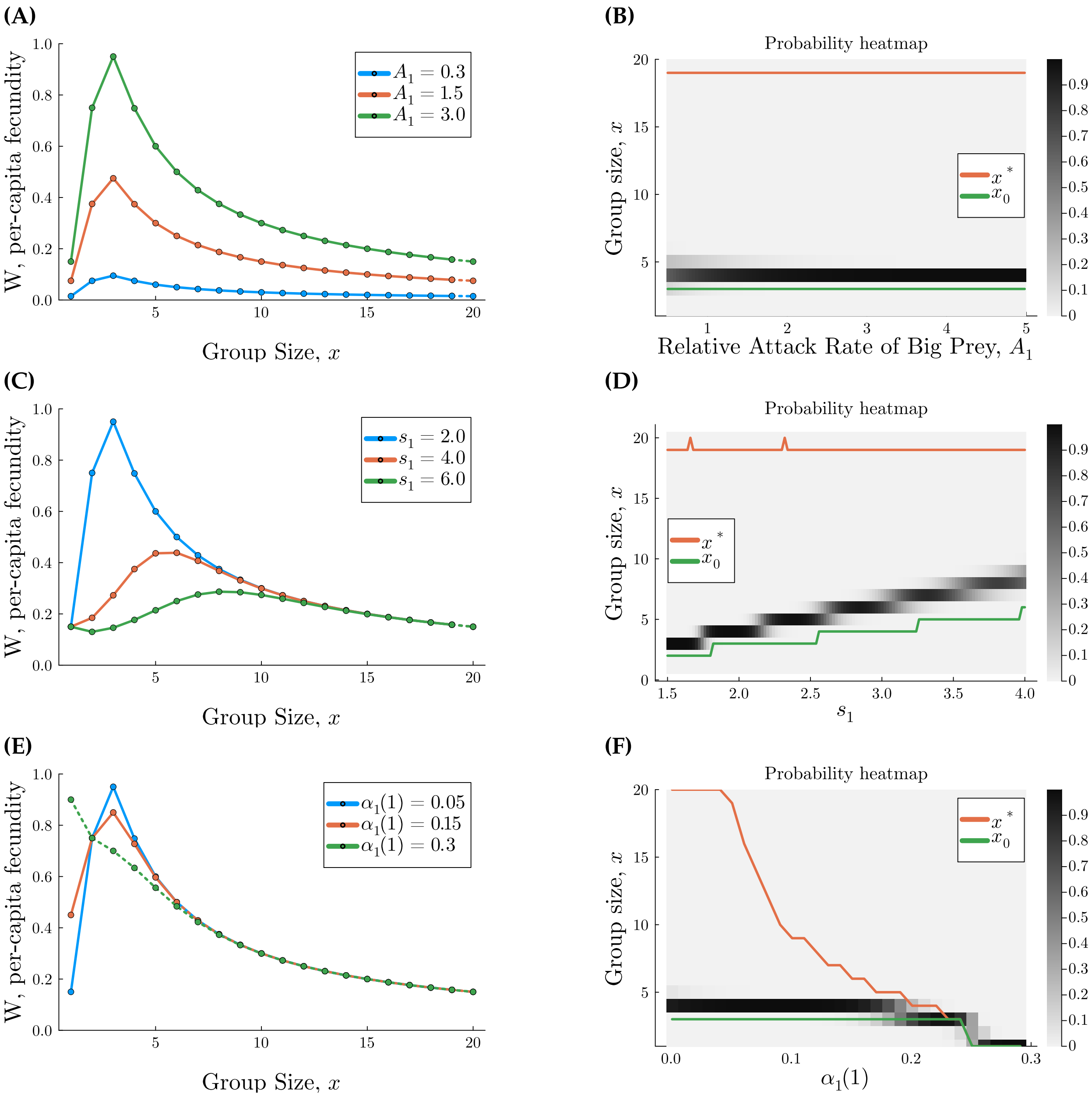


**Figure A.5:** The relationship between group size, fecundity, and the group size distribution at equilibrium without predator-prey population dynamics. Here we use a simple version of fecundity, $\tilde{W}(x)$ (eq. A.3). For all panels, $d = 100, P = 10$, and $x_m = 20$. All panels on the left side (**(A), (C),(E)**) show fecundity vs group size, with dotted lines indicating $\tilde{W}(x) < \tilde{W}(1)$. Panels on the right side (**(B), (D), (F)**) show the probability of experiencing a group of size $x$ at the stable equilibrium as heatmaps, with the curves indicating $x^*$, the Mangel and Clark predicted group size, and $x_0$, the group size that maximizes fecundity. Panels **(A), (B)** examine the effect of varying $A_1$, with $\alpha_1(1) = 0.05, s_1 = 2$. Panels **(C),(D)** examine the effect of varying the critical group size, $s_1$, for $A_1 = 3.0$ and $\alpha_1(1) = 0.05$. Panels **(E),(F)** examine the effect of varying $\alpha_1(1)$, for $s_1 = 2.0$ and $A_1 = 3.0$.

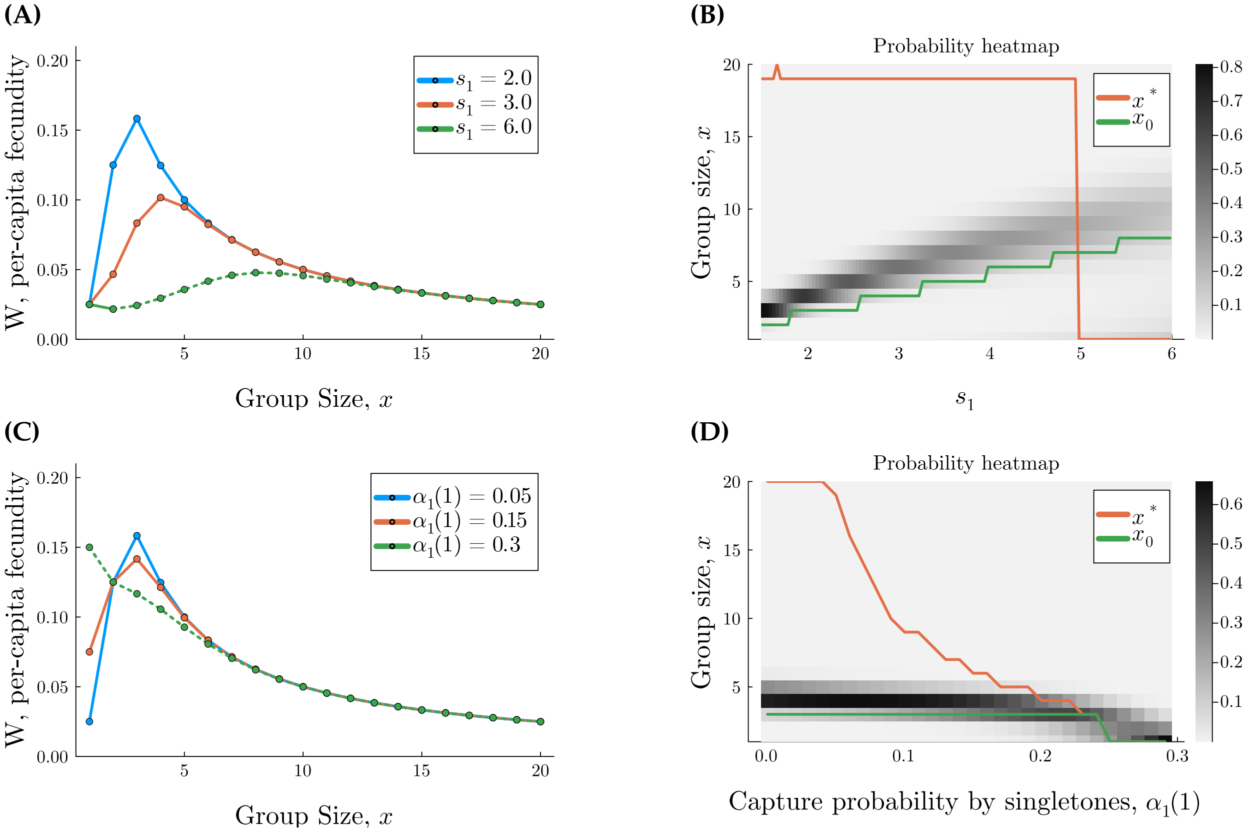


**Figure A.6:** The relationship between group size, fecundity, and the group size distribution at equilibrium, without predator or prey population dynamics. Here we use a simple version of fecundity, $\tilde{W}(x)$ (eq. A.3). For all panels, $d = 100, P = 10, A_1 = 0.5$ and $x_m = 20$. All panels on the left side (**(A), (C)**) show fecundity vs group size, with dotted lines indicating $\tilde{W}(x) < \tilde{W}(1)$. Panels on the right side (**(B), (D)**) show the probability of experiencing a group of size $x$, indicated with a heatmap (see colorbar), at the stable equilibrium. The orange curve shows the group size predicted by Clark and Mangel (1986), $x^*$, and the green curve shows the group size that maximizes fecundity, $x^*$. Panels **(A), (B)** examine the effect of varying the critical group size, $s_1$, for $\alpha_1(1) = 0.05$. Panels **(C),(D)** examine the effect of varying $\alpha_1(1)$, for $s_1 = 2.0$.

#### *A.2 Group Dynamics Equilibrium, No Population Dynamics*

Let $Q_1(\vec{g})$ be the right side of eq. 2a, $Q_2(\vec{g})$ be the right side of eq. 2b, $Q_{x_m}(\vec{g})$ be the right side of 2d, and let $Q_x(\vec{g})$ be the right side of 2c for any $x$ such that $x \in 3, 4, \cdots, x_m - 1$. Then from eqs. 2a-d,

$$\sum_{x=2}^{x_m} Q_x(\vec{g}) = -l \sum_{x=2}^{x_m} x g_x S(1,x) - g_1 \sum_{x=2}^{x_m-1} g_x S(x+1,1) + \frac{1}{2}\phi g_1^2 S(2,1) + l \sum_{x=3}^{x_m} x g_x S(1,x) + g_1 \sum_{x=2}^{x_m-1} g_x S(x+1,1),$$

which equals zero at equilibrium, so after simplifiying,

$$g_2 = \frac{1}{2}\phi g_1^2 \frac{S(2,1)}{2l(1 - S(2,1))}.$$

By setting $\sum_{y=x}^{x_m} Q_y(\vec{g}) = 0$, using similar logic, we find

$$g_x = g_1 g_{x-1}\left(\frac{S(x,1)}{lx(1 - S(x,1))}\right).$$

Let

$$c_x = \frac{S(x,1)}{lx(1 - S(x,1))}.$$

Then for $x = 2, 3, \cdots, x_m$, at equilibrium

$$g_x^* = \frac{1}{2}\phi g_1^{*x} c_x c_{x-1} \cdots c_2. \tag{A.4}$$

Note that even if population sizes are not constant (as is the case for the model where we meld group dynamics of predators with a one-predator, two-prey model), then we can separate timescales if the constant of group dynamics (after rescaling), $T_g$, is sufficiently small. We separate timescales by eliminating birth and death terms in order to find the equilibrium group size distribution $g_x^*$ for $N_1, N_2, P$ constant, and then substitute $g_x^*$ for $g_x$, for each $x$, into eqs. 12.

**Result 1.** *If population sizes are constant but group sizes change, then there is one unique equilibrium,*

$\mathbf{g}^*(x) = (g_1^*, g_2^*, \cdots, g_{x_m}^*)$, *as defined in eq. A.4*

*Proof.* To solve for $g_1$, we use the fact that $\sum_x xg_x = P$. Then solving for $P$ requires solving the polynomial of order $x_m$,

$$0 = -P + g_1 + \frac{1}{2}\phi\left(2g_1^2c(2) + 3g_1^3c(2)c(3) + ... + x_m g_1^{x_m}\prod_{y=2}^{x_m}c(y)\right)$$

The right side of the equation above is $-P$ if $g_1 = 0$ and increases as $g_1 \to \infty$, so it must have at least one positive root $g_1^*$.

Below, we use proof by contradiction to prove that there is only one positive root.

Say there is more than one positive root. Two of the roots are $\hat{g}_1, \tilde{g}_1$. If $g_1^* = \hat{g}_1$, then $g_x^* = \hat{g}_x = \frac{1}{2}\phi\hat{g}_1^x\prod_{y=2}^{x}c(y)$ from eq. A.4. If $g_1^* = \tilde{g}_1$, then $g_x^* = \tilde{g}_x = \frac{1}{2}\phi\tilde{g}_1^x\prod_{y=2}^{x}c(y)$. We must have $\sum_x xg_x = P$, so $\sum_x x\hat{g}_x = \sum_x x\tilde{g}_x$. Without loss of generality, say $\tilde{g}_1 > \hat{g}_1$. However, this means that for every positive integer $x$, $\tilde{g}_x > \hat{g}_x$, and thus $\sum_x x\hat{g}_x < \sum_x x\tilde{g}_x$, which is a contradiction. □

Note that division by $S(1,y)$ can cause issues for computation when $S(1,y)$ becomes, very, very small. To deal with this, we multiply both sides of the polynomial by $2\prod_{y=2}^{x_m} yS(1,y)$, i.e.,

$$0 = 2(g_1 - P)\prod_{y=2}^{x_m} yS(1,y) + \sum_{x=2}^{x_m} xg_1^x\left(\frac{1}{2}\phi\prod_{y=2}^{x}\frac{S(y,1)}{lyS(1,y)}\right)\left(2\prod_{y=2}^{x_m} yS(1,y)\right),$$

which simplifies to

$$0 = 2(g_1 - P)\prod_{y=2}^{x_m} yS(1,y) + \sum_{x=2}^{x_m-1}\frac{\phi}{l^{x-1}}xg_1^x\left(\prod_{y=2}^{x}S(y,1)\right)\left(\prod_{y=x+1}^{x_m} yS(1,y)\right) + \frac{\phi}{l^{x_m-1}}x_m g^{x_m}(1)\prod_{y=2}^{x_m}S(y,1).$$

#### *A.3 Jacobian of Group Dynamics without Population Dynamics*

Since $P$ is held constant, $g_{x_m} = P - g_1 - 2g_2 - \cdots - (x_m - 1)g_{x_m - 1}$. Let $\frac{\mathrm{d}g_x}{\mathrm{d}t} = Q_x(N_1, N_2, \vec{g})$ for $x = 1, 2, \cdots, x_m$, i.e., the right sides of eqs. 2 for $l = \phi = 1$. Since we are holding population dynamics constant, we remove the birth and death terms $x\tilde{W}(x)$ and $\tilde{\delta}$, and also simply set $T_g = 1$ since it does not effect local stability. Substituting for $g_{x_m}$ in terms of $g_1, g_2, \cdots, g_{x_m-1}$, $Q_1$ and

$Q_{x_m-1}$ become

$$Q_1(N_1, N_2, \vec{g}) = 2lg_2S(1,2) - \phi g_1^2 S(2,1) - x_m l S(1, x_m) g_1 + P l x_m S(1, x_m) + \sum_{x=2}^{x_m-1} g_x \left[ l x S(1,x) - g_1 S(x+1, 1) - l x x_m S(1, x_m) \right] \quad \text{(A.5)}$$

and

$$Q_{x_m-1}(N_1, N_2, \vec{g}) = -l(x_m - 1) g_{x_m-1} S(1, x_m - 1) - g_{x_m-1} g_1 S(x_m, 1) + g_{x_m-2} g_1 S(x_m - 1, 1) + l x_m S(1, x_m) \left[ P - \sum_{x=1}^{x_m-1} x g_x \right] \quad \text{(A.6)}$$

To find the Jacobian, we find the partial derivatives with respect to $g_1, g_2, ..., g_{x_m-1}$ of $Q_1, Q_2, \cdots, Q_{x_m-1}$. The first row of the Jacobian is

$$\frac{\partial Q_1}{\partial g_1} = -2lg_1 S(2,1) - x_m l S(1, x_m) - \sum_{x=2}^{x_m-1} g_x S(x+1, 1) \quad \text{(A.7)}$$

$$\frac{\partial Q_1}{\partial g_2} = 4lS(1,2) - g_1 S(3,1) - 2l x_m S(1, x_m) \quad \text{(A.8)}$$

$$\frac{\partial Q_1}{\partial g_x} = x l S(1,x) - g_1 S(x+1, 1) - x x_m l S(1, x_m) \quad \text{for } x > 2. \quad \text{(A.9)}$$

The second row is made up of

$$\frac{\partial Q_2}{\partial g_1} = -g_2 S(3,1) + \phi g_1 S(2,1) \quad \text{(A.10)}$$

$$\frac{\partial Q_2}{\partial g_2} = -2lS(1,2) - g_1 S(3,1) \quad \text{(A.11)}$$

$$\frac{\partial Q_2}{\partial g_3} = 3lS(1,3) \quad \text{(A.12)}$$

$$\frac{\partial Q_2}{\partial g_x} = 0 \quad \text{for } x > 3. \quad \text{(A.13)}$$

For rows $x = 3, 4, \cdots, x_m - 2$:

$$\frac{\partial Q_x}{\partial g_1} = -g_x S(x+1, 1) + g_{x-1} S(x, 1) \tag{A.14}$$
$$\frac{\partial Q_x}{\partial g_{x-1}} = g_1 S(x, 1) \tag{A.15}$$
$$\frac{\partial Q_x}{\partial g_x} = -xlS(1, x) - g_1 S(x+1, 1) \tag{A.16}$$
$$\frac{\partial Q_x}{\partial g_{x+1}} = l(x+1) S(1, x+1) \tag{A.17}$$
$$\frac{\partial Q_x}{\partial g_y} = 0 \quad \text{for } y \neq 1, x-1, x, x+1. \tag{A.18}$$

For row $x_m - 1$:

$$\frac{\partial Q_{x_m-1}}{\partial g_1} = -g_{x_m-1} S(x_m, 1) + g(x_m - 2) S(x_m - 1, 1) - x_m l S(1, x_m) \tag{A.19}$$
$$\frac{\partial Q_{x_m-1}}{\partial g_x} = -x x_m l S(1, x_m) \quad \text{for } x = 2, 3, \cdots, x_m - 3 \tag{A.20}$$
$$\frac{\partial Q_{x_m-1}}{\partial g_{x_m-2}} = g_1 S(x_m - 1, 1) - l(x_m - 2) x_m S(1, x_m) \tag{A.21}$$
$$\frac{\partial Q_{x_m-1}}{\partial g_{x_m-1}} = -l(x_m - 1) S(1, x_m - 1) - g_1 S(x_m, 1) - l(x_m - 1) x_m S(1, x_m) \tag{A.22}$$

#### B Scaling Population Parameters using Scaling Laws

Following the scaling relationships used in the model from Weitz and Levin (2006), we assume the basal metabolic rate of prey type $i$ is proportional to $m^{\epsilon}$, where $2/3 < \epsilon < 1$ (and for most taxa it tends to be 3/4), so that carrying capacity $k_i \propto m_i^{-\epsilon}$ and the intrinsic growth rate $r_i \propto m_i^{\epsilon-1}$. Then $k_1/k_2 = (m_1/m_2)^{-\epsilon}$ and $r_1/r_2 = (m_1/m_2)^{1-\epsilon}$. For $m_p$ the mass of predators, the conversion of prey eaten to predators produced is proportional to $m_i/m_p$, so $b_1/b_2 = m_1/m_2$. If the population size of big prey is infinitely large, i.e., $M_i \to \infty$, then the growth of singleton predators becomes $\frac{b_i}{h_i(1)}$. Following Weitz and Levin (2006), the scaling should be like that of prey growing with infinite resources, so

$$\frac{b_i}{h_i(1)} \propto m_p^{\theta-1}$$

for $m_p$ the mass of predators and $\theta$ a positive constant. Then $h_1(1)/h_2(1) = b_1/b_2$.

#### C Non-dimensionalization of Population Dynamics Model

The non-dimensionalized functional response is

$$\tilde{f}_i(x, N_1, N_2) = \frac{A_i \alpha_i(x) N_i}{1 + H_1(x)\alpha_1(x)N_1 + H_2(x)\alpha_2 N_2} = \frac{\frac{a_i}{\nu}\alpha_i(x)\frac{M_i}{k_i}}{1 + a_1 h_1(x)\alpha_1(x)M_1 + a_2 h_2(x)\alpha_2(x)M_2}$$
$$= \frac{f_i(x, M_1, M_2)}{k_i \nu}. \quad \text{(C.23)}$$

Let the rescaled fecundity be $\tilde{W}$, where we define

$$\tilde{W}(x) = \frac{1}{x}\left(\beta_1 \tilde{f}_1(x, N_1, N_2) + \beta_2 \tilde{f}_2(x, N_1, N_2)\right). \quad \text{(C.24)}$$

Then substituting from eq. C.23,

$$\tilde{W}(x) = \frac{1}{x}\left(\frac{\beta_1}{k_1\nu} f_1(x, M_1, M_2) + \frac{\beta_2}{k_2\nu} f_1(x, M_1, M_2)\right).$$

Substiting the definition $\beta_i = b_i k_i$ (see Table 2), this means that $\tilde{W}(x) = W(x)/\nu$. The best response function is

$$S(x, y) = \frac{1}{1 + \exp\left[-d\left(\tilde{W}(x) - \tilde{W}(1)\right)\right]} \tag{C.25}$$

for $d = \gamma\nu$.

### D Group Formation Decisions

Here, we explore the conditions under which predators should join groups, if given the chance, in response to prey availability. Fecundity is derived from the benefits of hunting i.e., $\tilde{W}$ is defined (after non-dimensionalization) in eq. 13. Since fitness is determined by functional responses, we determine the effect of group size on the functional responses to big prey and small prey. The functional response to big prey increases with group size and the per-capita functional response to small prey, $\tilde{f}_2(x)/x$, decreases with group size, as shown below in Results 2 and 3.

**Result 2.** *The functional response to big prey increases with group size, i.e.,* $\tilde{f}_1(x+1, N_1, N_2) > \tilde{f}_1(x, N_1, N_2)$.

*Proof.* This result is true if

$$\frac{\alpha_1(x+1)N_1}{1 + \alpha_1(x+1)H_1(x+1)N_1 + \alpha_2 H_2(x+1)N_2} > \frac{\alpha_1(x)N_1}{1 + \alpha_1(x)H_1(x)N_1 + \alpha_2 H_2(x)N_2}$$

which is rearranged to be

$$\underbrace{\alpha_1(x+1) - \alpha_1(x)}_{I} + \underbrace{N_1\alpha_1(x)\alpha_1(x+1)\left[H_1(x) - H_1(x+1)\right]}_{II} + \underbrace{N_2\alpha_2\left(\alpha_1(x+1)H_2(x) - \alpha_1(x)H_2(x+1)\right)}_{III} > 0. \tag{D.26}$$

This is true because terms I, II, and III are positive, since we define $H_i(x) = H_{ia} + \frac{1}{x}H_{ib}$.

□

**Result 3.** *The per capita functional response to small prey decreases as x increases, i.e.,* $\tilde{f}_2(1, N_1, N_2) > \frac{1}{2}\tilde{f}_2(2, N_1, N_2) > \cdots > \frac{1}{x_m}\tilde{f}_2(x_m, N_1, N_2)$

*Proof.* To show that the per capita functional response to small prey decreases as $x$ increases, from the definition of the functional response in eq. 4, we want to show that

$$(x+1)\left[1+\alpha_1(x+1)H_1(x+1)N_1+\alpha_2 H_2(x+1)N_2\right]-x\left[1+\alpha_1(x)H_1(x)N_1+\alpha_2 H_2(x)N_2\right]>0$$

for all $x \geq 1$. The left side of the inequality above is equal to

$$1+N_1\left[(x+1)H_1(x+1)\alpha_1(x+1)-xH_1(x)\alpha_1(x)\right]+N_2\left[(x+1)H_2(x+1)-xH_2(x)\right],$$

and by the definitions of $H_i(x)$ and $\alpha_1(x)$, $(x+1)H_i(x+1) > xH_i(x)$ and $\alpha_1(x+1) > \alpha_1(x)$, so the above expression is positive. □

This means that predation pressure on small prey is highest if all predators are solitary. Under some conditions, the functional response of a group to small prey can be shown to also decrease with group size. This is explored below.

**Result 4.** *If handling time does not decrease with group size (i.e., $H_1 = H_{1a}$, $H_2 = H_{2a}$) then the functional response to small prey decreases with group size.*

*Proof.* Showing the functional response to small prey (defined in eq. 4) decreases with group size if $H_i = H_{ia}$ for $i = 1, 2$ requires showing that

$$1+\alpha_1(x+1)H_1(x+1)N_1+\alpha_2 H_2(x+1)N_2 > 1+\alpha_1(x)H_1(x)N_1+\alpha_2 H_2(x)N_2,$$

or equivalently (using $H_i = H_{ia}$ for brevity),

$$H_1 N_1\left[\alpha_1(x+1)-\alpha_1(x)\right]>0,$$

which is true by the definition of $\alpha_1(x)$. □

Groups should form in a population of solitary predators if the fecundity of being in a group of size 2 is greater than that of being in a group of size 1. The conditions for this are presented in the next result.

**Result 5.** *If the handling times and benefits of big prey and small prey are scaled by bodymass, such that* $H_1(1) = \left(\frac{A_1\beta_1}{A_2\beta_2}\right) H_2(1)$*, then the fecundity of being in a group of size* $x$ *is greater than being alone, i.e.,* $\tilde{W}(x) > \tilde{W}(1)$*, if*

$$\frac{\beta_1}{\beta_2} > \left(\frac{A_2\alpha_2 N_2}{A_1 N_1}\right)\left(\frac{(x-1)\left[1+\alpha_1(x)N_1H_{1a}+\alpha_2N_2H_{2a}\right]}{\alpha_1(x)-x\alpha_1(1)-\alpha_1(1)(x-1)\left(\alpha_1(x)N_1H_{1a}+\alpha_2N_2H_{2a}\right)}\right). \tag{D.27}$$

*If* $H_{2a} = H_{1a} = 0$*, then* $\tilde{W}(x) > \tilde{W}(1)$ *if* $\alpha_1(x) - x\alpha_1(1) > 0$ *and*

$$\frac{\beta_1}{\beta_2} > \frac{\alpha_2 A_2 N_2 \,(x-1)}{A_1 N_1\left[\alpha_1(x) - x\alpha_1(1)\right]}. \tag{D.28}$$

*Proof.* From eq. 10, after non-dimensionalizing, the fecundity of being in a group of size $x$ is greater than being alone if

$$\frac{1}{x}\left(\beta_1\tilde{f}_1(x)+\beta_2\tilde{f}_2(x)\right) > \beta_1\tilde{f}_1(1)+\beta_2\tilde{f}_2(1),$$

or equivalently,

$$\beta_1\left(\frac{1}{x}\tilde{f}_1(x)-\tilde{f}_1(1)\right) > \beta_2\left(\tilde{f}_2(1)-\frac{1}{x}\tilde{f}_2(x)\right). \tag{D.29}$$

If $\frac{1}{x}\tilde{f}_1(x) - \tilde{f}_1(1) > 0$, then inequality D.29 is true if

$$\frac{\beta_1}{\beta_2} > \frac{\tilde{f}_2(1)-\frac{1}{x}\tilde{f}_2(x)}{\frac{1}{x}\tilde{f}_1(x)-\tilde{f}_1(1)}. \tag{D.30}$$

and if $\frac{1}{x}\tilde{f}_1(x) - \tilde{f}_1(1) < 0$, inequality D.29 is true if

$$\frac{\beta_1}{\beta_2} < \frac{\tilde{f}_2(1)-\frac{1}{x}\tilde{f}_2(x)}{\frac{1}{x}\tilde{f}_1(x)-\tilde{f}_1(1)}.$$

However, from Result 3, $\tilde{f}_2(1) - \frac{1}{x}\tilde{f}_2(x) > 0$ because $\tilde{f}_2(1) > \tilde{f}_2(x)$, so the fraction above is negative. By definition, $\beta_1, \beta_2 > 0$, so the per capita fecundity of being in a group of size $x$ is greater than being alone if inequality D.30 is true.

Let $B = \beta_1/\beta_2$ and let $A = A_1/A_2$ for brevity in the following calculations. From inequality

D.29, a group of size $x$ is favored over being alone if,

$$AB\left[\frac{1}{x}\left(\frac{N_1\alpha_1(x)}{1+\alpha_1(x)H_1(x)N_1+\alpha_2H_2(x)N_2}\right)-\frac{N_1\alpha_1(1)}{1+\alpha_1(1)H_1(1)N_1+\alpha_2H_2(1)N_2}\right] \\ -\left[\frac{N_2\alpha_2}{1+\alpha_1(1)H_1(1)N_1+\alpha_2H_2(1)N_2}-\frac{1}{x}\left(\frac{N_2\alpha_2}{1+\alpha_1(x)H_1(x)N_1+\alpha_2H_2(x)N_2}\right)\right]>0, \quad \text{(D.31)}$$

or equivalently,

$$AB\left\{N_1\alpha_1(x)\left[1+\alpha_1(1)H_1(1)N_1+\alpha_2H_2(1)N_2\right]-xN_1\alpha_1(1)\left[1+\alpha_1(x)H_1(x)N_1+\alpha_2H_2(x)N_2\right]\right\} \\ -\left\{xN_2\alpha_2\left[1+\alpha_1(x)H_1(x)N_1+\alpha_2H_2(x)N_2\right]-N_2\alpha_2\left[1+\alpha_1(1)H_1(1)N_1+\alpha_2H_2(1)N_2\right]\right\}>0. \quad \text{(D.32)}$$

Let $w_{1x}=\alpha_1(x)N_i$, $w_2=\alpha_2N_2$ for brevity. After substituting, the inequality in eq. D.31 is equivalent to

$$AB\left\{w_{1x}\left[1+w_{11}H_1(1)+w_2H_2(1)\right]-xw_{11}\left[1+w_{1x}H_1(x)+w_2H_2(x)\right]\right\} \\ -\left\{xw_2\left[1+w_{1x}H_1x)+w_2H_2(x)\right]-w_2\left[1+w_{11}H_1(1)+w_2H_2(1)\right]\right\}>0. \quad \text{(D.33)}$$

With the scaling assumptions, $H_1(1)=ABH_2(1)$ and $H_2(1)=H_1(1)/(AB)$, so after substituting strategically,

$$AB\left\{w_{1x}\left[1+w_{11}H_1(1)\right]-xw_{11}\left[1+w_{1x}H_1(x)+w_2H_2(x)\right]\right\}+w_2w_{1x}H_1(1) \\ -\left\{xw_2\left[1+w_{1x}H_1x)+w_2H_2(x)\right]-w_2\left[1+w_2H_2(1)\right]\right\}+ABw_2w_{11}H_2(1)>0. \quad \text{(D.34)}$$

After regrouping,

$$AB\left\{w_{1x}\left[1+w_{11}H_1(1)\right]-xw_{11}\left[1+w_{1x}H_1(x)+w_2H_2(x)\right]+w_2w_{11}H_2(1)\right\} \\ -\left\{xw_2\left[1+w_{1x}H_1(x)+w_2H_2(x)\right]-w_2\left[1+w_2H_2(1)\right]-w_2w_{1x}H_1(1)\right\}>0. \quad \text{(D.35)}$$

Now distribute and regroup:

$$AB\left\{w_{1x}-xw_{11}+w_{1x}w_{11}\left(H_1(1)-xH_1(x)\right)+w_{11}w_2\left(H_2(1)-xH_2(x)\right)\right\} \\ -\left\{xw_2-w_2+w_2w_{1x}\left(xH_1(x)-H_1(1)\right)+w_2^2\left(xH_2(x)-H_2(1)\right)\right\}>0. \quad \text{(D.36)}$$

Since $H_i(x) = H_{ia} + H_{ib}/x$, $xH_1(x) - H_1(1) = H_{1a}(x-1)$ and $xH_2(x) - H_2(x) = H_{2a}(x-1)$. Then we have

$$AB\left\{w_{1x} - xw_{11} + w_{11}(1-x)\left(w_{1x}H_{1a} + w_2H_{2a}\right)\right\} - w_2(x-1)\left(1 + w_{1x}H_{1a} + w_2H_{2a}\right) > 0.$$

Isolating $B$,

$$B > \left(\frac{A_2}{A_1}w_2\right)\left(\frac{(x-1)\left(1 + w_{1x}H_{1a} + w_2H_{2a}\right)}{w_{1x} - xw_{11} + w_{11}(1-x)\left(w_{1x}H_{1a} + w_2H_{2a}\right)}\right). \quad \text{(D.37)}$$

Now substitute the definitions of $w_{1x}, w_2$, and $B$:

$$\frac{\beta_1}{\beta_2} > \left(\frac{A_2N_2\alpha_2}{A_1N_1}\right)\left(\frac{(x-1)\left(1 + N_1\alpha_1(x)H_{1a} + \alpha_2N_2H_{2a}\right)}{\alpha_1(x) - x\alpha_1(1) + \alpha_1(1)(1-x)\left(\alpha_1(x)N_1H_{1a} + \alpha_2N_2H_{2a}\right)}\right) \quad \text{(D.38)}$$

If $H_{1a} = H_{2a} = 0$, then groups of size $x$ are favored over being solitary, i.e., $\tilde{W}(x) > \tilde{W}(1)$, if

$$\frac{\beta_1}{\beta_2} > \frac{\alpha_2A_2N_2\,(x-1)}{A_1N_1\left[\alpha_1(x) - x\alpha_1(1)\right]}.$$

□

If the benefit of big prey is higher — which could happen if the body size of big prey is very large — then a lower population size of big prey is required for groups to be favored. The minimum population sizes and benefits of big prey required such that $\tilde{W}(x) > \tilde{W}(1)$ generally increase with $x$ (Fig. D.7). For the parameters in Fig. D.7, an exception to this trend occurs for groups of size 2 and 3, which have indistinguishable threshold curves for $s = 2$. However, if the critical group size $s$ is slightly increased to $s = 2.45$, groups of 3 are favored over being alone for smaller prey abundances and benefit ratios than are required for pairs to be favored. Thus increasing the critical group size, which corresponds to the inflection point of the capture probability curve, can lead to regions where larger groups could be favored over being solitary, but these groups do not form because it is not favorable to form pairs.

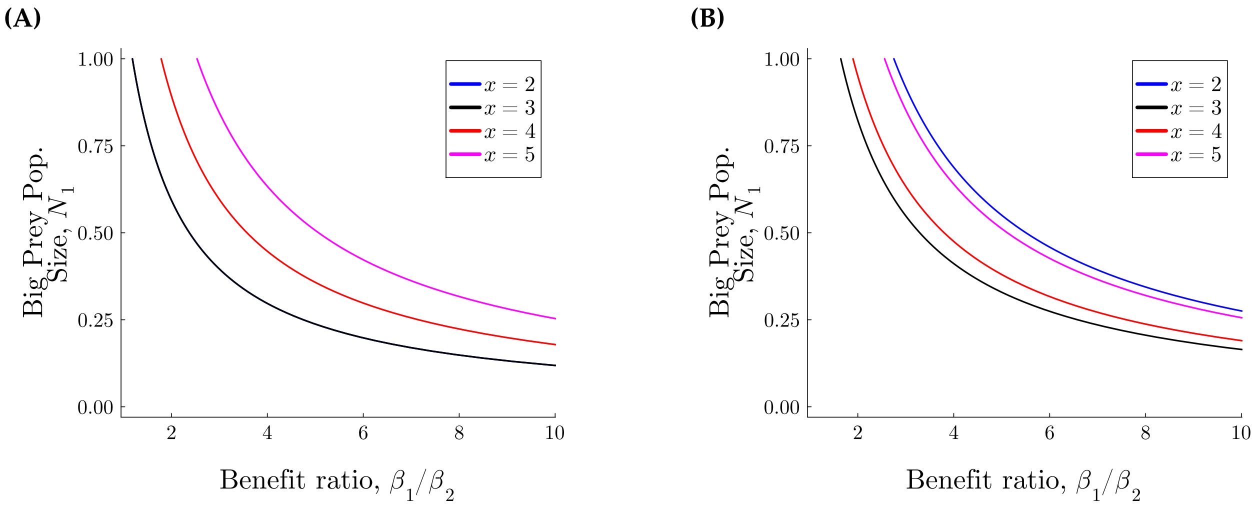


**Figure D.7:** Prey characteristics at which groups should form, as determined by Result **??**, if **(A)** $s_1 = 2$, and **(B)** $s_1 = 2.45$. If the ratio $\beta_1/\beta_2$ is greater than a curve, the per-capita fitness of being in a group of the corresponding group size ($x = 1, 2, 3, 4$, or 5) is greater than that of being alone. Here $A_1\beta_1/(A_2\beta_2) = H_{1b}/H_{2b}$, the scaled population size of small prey is $N_2 = 0.5$, and the parameters are $H_{2b} = 1$, $\alpha_2 = 0.95$, and $\alpha_1(1) = 0.05$. There is no group-size independent component of handling time, i.e., $H_{1a} = H_{2a} = 0$

### E Population Dynamics without Group Dynamics

If all predators are in a group of size $x = x^*$, then one-predator two-prey dynamics become

$$\text{(predator)} \quad \frac{\mathrm{d}P}{\mathrm{d}T} = P\left[\frac{\beta_1 A_1 \alpha_1(x) N_1 + \beta_2 A_2 \alpha_2}{x\left(1 + \alpha_1(x) H_1(x) N_1 + \alpha_2 H_2(x) N_2\right)} - \tilde{\delta}\right] \tag{E.39a}$$

$$\text{(prey } i\text{)} \quad \frac{\mathrm{d}N_i}{\mathrm{d}T} = \eta_i N_i \left(1 - N_i\right) - \frac{A_i \alpha_i(x) P N_i}{x\left(1 + \alpha_1(x) H_1(x) N_1 + \alpha_2 H_2(x) N_2\right)} \tag{E.39b}$$

The nullclines are $P = 0$, $N_1 = 0$, $N_2 = 0$, or

$$\text{(predator)} \quad x\tilde{\delta} = \frac{\beta_1 A_1 \alpha_1(x) N_1 + \beta_2 A_2 \alpha_2 N_2}{1 + \alpha_1(x) H_1(x) N_1 + \alpha_2 H_2(x) N_2} \tag{E.40a}$$

$$\text{(prey } i\text{)} \quad \eta_i(1 - N_i) = \frac{A_i \alpha_i(x) P}{x\left(1 + \alpha_1(x) H_1(x) N_1 + \alpha_2 H_2(x) N_2\right)} \tag{E.40b}$$

respectively. From the predator nullcline, eq. E.40a, at the coexistence equilibrium,

$$N_1^* = c_1(x) N_2 + c_0(x) \tag{E.41}$$

where,

$$c_0(x) = \frac{\tilde{\delta}x}{\alpha_1(x)\left(\beta_1 A_1 - H_1(x)\tilde{\delta}x\right)}, \text{ and } c_1(x) = \frac{\alpha_2\left(H_2(x)\tilde{\delta}x - \beta_2 A_2\right)}{\alpha_1(x)\left(\beta_1 A_1 - H_1(x)\tilde{\delta}x\right)}.$$

Note that with scaling assumptions as expressed in eq. 8, **if** $H_{1a} = H_{2a} = 0$ **(so the group-independent components of handling time are neglected)** then $\frac{\beta_2 A_2}{H_2(x)} = \frac{\beta_1 A_1}{H_1(x)}$ so $c_1(x) = -\frac{\alpha_2 H_2(x)}{\alpha_1(x)H_1(x)}$, and for $N_2^*$ to be positive, $\frac{\beta_1 A_1}{H_2(x)} > \tilde{\delta}x$.

From the prey nulllcines (eq. E.40b for $i = 1, 2$):

$$A_1\alpha_1(x)P/x = \eta_1(1 - N_1)\left(1 + \alpha_1(x)H_1(x)N_1 + \alpha_2 H_2(x)N_2\right),$$

$$\text{and } A_2\alpha_2 P/x = \eta_2(1 - N_2)\left(1 + \alpha_1(x)H_1(x)N_1 + \alpha_2 H_2(x)N_2\right),$$

so

$$\frac{A_1\alpha_1(x)}{\eta_1(1 - N_1)} = \frac{A_2\alpha_2}{\eta_2(1 - N_2)}.$$

Substituting eq. E.41 for $N_1$,

$$N_2^* = \frac{\gamma_2(x)(1 - c_0(x)) - \gamma_1(x)}{\gamma_2(x)c_1(x) - \gamma_1} \quad \text{where} \quad \gamma_i(x) = \frac{A_i\alpha_i(x)}{\eta_i} \tag{E.42}$$

We can substitute eq. E.42 into E.41 to find $N_1^*$. Finally, both can be substituted into the eq. E.40b to find $P^*$.

If big prey is absent, but the other populations persist, then

$$N_2^* = \frac{x\tilde{\delta}}{\alpha_2\left(\beta_2 A_2 - xH_2(x)\tilde{\delta}\right)}, \quad P^* = \left(\frac{\eta_2\beta_2}{\tilde{\delta}}\right) N_2^*(1 - N_2^*)$$

Since $0 < N_2 \leq 1$, this equilibrium exists if $x\tilde{\delta} \leq \alpha_2(\beta_2 A_2 - xH_2(x)\tilde{\delta})$ and $\beta_2 A_2 > xH_2(x)\tilde{\delta}$. Simplifying, that means we have an upper bound on $x$ for predators and small prey to persist without big prey:

$$x \leq \frac{\alpha_2(\beta_2 A_2 - H_{2b}\tilde{\delta})}{\tilde{\delta}(1 + \alpha_2 H_{2a})}.$$

That means that predators and small prey can coexist without big prey for predators in large groups if both group-independent and dependent portions of the handling time are small, the scaled death rate is small, and the scaled benefit and attack rate on small prey are large. Also note that $0 < N_2 \leq 1$ means that $P^* \leq \frac{\eta_2 \beta_2}{4\tilde{\delta}}$.

On the other hand, if small prey is absent, i.e. $N_2^* = 0$, but the other populations persist, then

$$N_1^* = \frac{x\tilde{\delta}}{\alpha_1(x)\left(\beta_1 A_1 - xH_1(x)\tilde{\delta}\right)}, \quad P^* = \left(\frac{\eta_1\beta_1}{\tilde{\delta}}\right) N_1^*(1 - N_1^*). \tag{E.43}$$

Since $0 < N_1 \leq 1$, this equilibrium exists if $x\tilde{\delta} \leq \alpha_1(x)(\beta_1 A_1 - xH_1(x)\tilde{\delta})$ and $\beta_1 A_1 > xH_1(x)\tilde{\delta}$. Due to the definition of $\alpha_1(x)$, we cannot isolate bounds on $x$.

#### *E.1 Local Stability, Population Dynamics without Group Dynamics*

The Jacobian for both prey types present is,

$$\Omega = \begin{pmatrix} -\tilde{\delta} + \tilde{W}(x) & P\frac{\partial \tilde{W}}{\partial N_1} & P\frac{\partial \tilde{W}}{\partial N_2} \\ -\frac{1}{x}\tilde{f}_1 & \eta_1(1 - 2N_1) - \frac{P}{x}\frac{\partial \tilde{f}_1}{\partial N_1} & -\frac{P}{x}\frac{\partial \tilde{f}_1}{\partial N_2} \\ -\frac{1}{x}\tilde{f}_2 & -\frac{P}{x}\frac{\partial \tilde{f}_2}{\partial N_1} & \eta_2(1 - 2N_2) - \frac{P}{x}\frac{\partial \tilde{f}_2}{\partial N_2} \end{pmatrix}, \tag{E.44}$$

where we suppress the dependence of $\tilde{W}, f_i$ on $x, N_1, N_2$ for brevity in the Jacobian above. Note that,

$$\frac{\partial \tilde{f}_i}{\partial N_i} = \frac{A_i \alpha_i(x)\left(1 + \alpha_j(x) H_j(x) N_j\right)}{\left(1 + \alpha_1(x)H_1(x)N_1 + \alpha_2 H_2(x) N_2\right)^2} \quad \text{and} \quad \frac{\partial \tilde{f}_i}{\partial N_j} = -\frac{A_i \alpha_1(x)\alpha_2 H_j(x) N_i}{\left(1 + \alpha_1(x)H_1(x)N_1 + \alpha_2 H_2(x) N_2\right)^2}, \tag{E.45}$$

for $i, j = 1, 2$. As for fecundity,

$$\begin{aligned} \frac{\partial \tilde{W}}{\partial N_i} &= \frac{1}{x}\left(\beta_i \frac{\partial \tilde{f}_i}{\partial N_i} + \beta_j \frac{\partial \tilde{f}_j}{\partial N_i}\right) \\ &= \frac{1}{x}\left(\frac{\beta_i A_i \alpha_i(x) + \alpha_1(x)\alpha_2 N_j\left(\beta_i A_i H_j(x) - \beta_j A_j H_i(x)\right)}{\left(1 + \alpha_1(x)H_1(x)N_1 + \alpha_2 H_2(x) N_2\right)^2}\right), \end{aligned} \tag{E.46}$$

and if the parameters are scaled by bodymass (see eq. 6), then $\beta_i A_i H_j(x) - \beta_j A_j H_i(x) = 0$, so $\frac{\partial \tilde{W}}{\partial N_i} > 0$. The characteristic polynomial is

$$0 = \lambda^3 - \mathrm{Tr}(\Omega)\lambda^2 + \lambda\left(\Omega_{12}\Omega_{21} + \Omega_{13}\Omega_{31} + \Omega_{23}\Omega_{32} - \Omega_{11}\Omega_{22} - \Omega_{11}\Omega_{33} - \Omega_{22}\Omega_{33}\right) - \det(\Omega).$$

At the coexistence equilibrium, the jacobian is

$$\Omega = \begin{pmatrix} 0 & P\frac{\partial \tilde{W}}{\partial N_1} & P\frac{\partial \tilde{W}}{\partial N_2} \\ \frac{1}{x}f_1 & \eta_1(1-2N_1) - \frac{P}{x}\frac{\partial \tilde{f}_1}{\partial N_1} & -\frac{P}{x}\frac{\partial f_1}{\partial N_2} \\ -\frac{1}{x}f_2 & -\frac{P}{x}\frac{\partial f_2}{\partial N_1} & \eta_2(1-2N_2) - \frac{P}{x}\frac{\partial \tilde{f}_2}{\partial N_2} \end{pmatrix},$$

If big prey is extinct but predators and small prey are not, $\frac{\partial \tilde{f}_1}{\partial N_2} = \tilde{f}_1 = 0$ and $\beta_2 \tilde{f}_2 / x - \tilde{\delta} = 0$, so

$$\Omega = \begin{pmatrix} 0 & P\frac{\partial \tilde{W}}{\partial N_1} & P\frac{\partial \tilde{W}}{\partial N_2} \\ 0 & \eta_1 - \frac{P}{x}\frac{\partial \tilde{f}_1}{\partial N_1} & 0 \\ -\frac{1}{x}\tilde{f}_2 & -\frac{P}{x}\frac{\partial \tilde{f}_2}{\partial N_1} & \eta_2(1-2N_2) - \frac{P}{x}\frac{\partial \tilde{f}_2}{\partial N_2}, \end{pmatrix}$$

for which the eigenvalues are

$$\lambda_1 = \eta_1 - \frac{P}{x}\left(\frac{\partial f_1}{\partial N_1}\right)$$

and $\lambda_2, \lambda_3$ are the solutions to the quadratic equation

$$0 = \lambda^2 - \lambda\left[\eta_2(1-2N_2) - \frac{P}{x}\frac{\partial \tilde{f}_2}{\partial N_2}\right] + \beta_2 \frac{P f_2}{x^2}\frac{\partial f_2}{\partial N_2}.$$

Since the quadratic is positive at $\lambda = 0$ and the leading coefficient is positive, its roots must be complex. The real part of the eigenvalues is $\eta_2(1-2N_2) - \frac{P}{x}\frac{\partial \tilde{f}_2}{\partial N_2}$.

If predators are extinct, then the jacobian is,

$$\Omega = \begin{pmatrix} -\tilde{\delta} + \tilde{W}(x) & 0 & 0 \\ -\tilde{f}_1/x & \eta_1(1-2N_1) & 0 \\ -\tilde{f}_2/x & 0 & \eta_2(1-2N_2) \end{pmatrix}.$$

Note that $N_1 = N_2 = 1$ if predators are extinct, so this equilibrium is unstable if predators in groups of size $x$ can survive with the maximal amount of prey available.

##### *E.1.1 Local Stability, One Prey Population*

The Jacobian for only big prey present is

$$\Omega = \begin{pmatrix} -\tilde{\delta} + \tilde{W}(x) & P\frac{\mathrm{d}\tilde{W}}{\mathrm{d}N_1} \\ -\tilde{f}_1/x & \eta_1(1-2N_1) - \frac{P}{x}\frac{\mathrm{d}\tilde{f}_1}{\mathrm{d}N_1} \end{pmatrix}, \tag{E.47}$$

where $\frac{\mathrm{d}\tilde{W}}{\mathrm{d}N_1} = \frac{1}{x}\beta_1\frac{\mathrm{d}\tilde{f}_1}{\mathrm{d}N_1}$, $\tilde{f}_1 = \tilde{f}_1(x, N_1)$, and

$$\frac{\mathrm{d}\tilde{f}_1}{\mathrm{d}N_1} = \frac{A_1\alpha_1(x)}{(1+\alpha_1(x)H_1(x)N_1)^2}.$$

For $\lambda$ an eigenvalue, the characteristic polynomial of $\Omega$ is

$$\mathrm{Char}(\lambda) = \lambda^2 - \lambda\left[-\tilde{\delta} + \tilde{W}(x) + \eta_1(1-2N_1) - \frac{P}{x}\frac{\mathrm{d}\tilde{f}_1}{\mathrm{d}N_1}\right] + \left(\tilde{W}(x) - \tilde{\delta}\right)\left[\eta_1(1-2N_1) - \frac{P}{x}\frac{\mathrm{d}\tilde{f}_1}{\mathrm{d}N_1}\right] + \frac{1}{x}\tilde{f}_1 P\frac{\mathrm{d}\tilde{W}(x)}{\mathrm{d}N_1},$$

and since $\frac{\mathrm{d}\tilde{W}(x)}{\mathrm{d}N_1}\frac{1}{x}\beta_1\frac{\mathrm{d}\tilde{f}_1}{\mathrm{d}N_1}$ and $\tilde{W}(x) = \frac{1}{x}\beta_1\tilde{f}_1$, the characteristic polynomial can be simplified to

$$\mathrm{Char}(\lambda) = \lambda^2 - \lambda\left[\frac{\beta_1}{x}\tilde{f}_1 - \tilde{\delta} + \eta_1(1-2N_1) - \frac{1}{x}P\frac{\mathrm{d}\tilde{f}_1}{\mathrm{d}N_1}\right] + \eta_1\left(\frac{1}{x}\beta_1\tilde{f}_1 - \tilde{\delta}\right)(1-2N_1) + \frac{1}{x}\tilde{\delta}P\frac{\mathrm{d}\tilde{f}_1}{\mathrm{d}N_1}.$$

At the coexistence equilibrium, $\tilde{W}_1 - \tilde{\delta} = 0$ and so the eigenvalues are

$$\lambda = \frac{1}{2}\left\{\underbrace{\eta_1(1-2N_1) - \frac{1}{x}P\frac{\mathrm{d}\tilde{f}_1}{\mathrm{d}N_1}}_{\mathrm{I}} \pm \underbrace{\sqrt{\left[\eta_1(1-2N_1) - \frac{P}{x}\frac{\mathrm{d}\tilde{f}_1}{\mathrm{d}N_1}\right]^2 - \frac{4}{x^2}P\beta_1\tilde{f}_1\frac{\mathrm{d}\tilde{f}_1}{\mathrm{d}N_1}}}_{\mathrm{II}}\right\}$$

The magnitude of term II is at most the magnitude of term I if term II is real. If term I is positive, the eigenvalues are positive and any equilibrium is unstable, whereas if term I is negative, the eigenvalues are negative and any equilibrium is stable. Thus a coexistence equilibrium is stable if

$$\eta_1(1-2N_1) - \frac{P}{x}\frac{\mathrm{d}\tilde{f}}{\mathrm{d}N_1} < 0.$$

If predators are extinct, the Jacobian becomes

$$\Omega = \begin{pmatrix} \tilde{W} - \tilde{\delta} & 0 \\ -\tilde{f}_1/x & -\eta_1, \end{pmatrix}$$

so the eigenvalues are $\tilde{W} - \tilde{\delta}$ and $-\eta_1$, and the equilibrium is a saddle if $\tilde{W} > \tilde{\delta}$ and stable otherwise.

#### E.2 Plots of Stable Equilibria, varying Scale and Group Size

##### E.2.1 One Prey Population

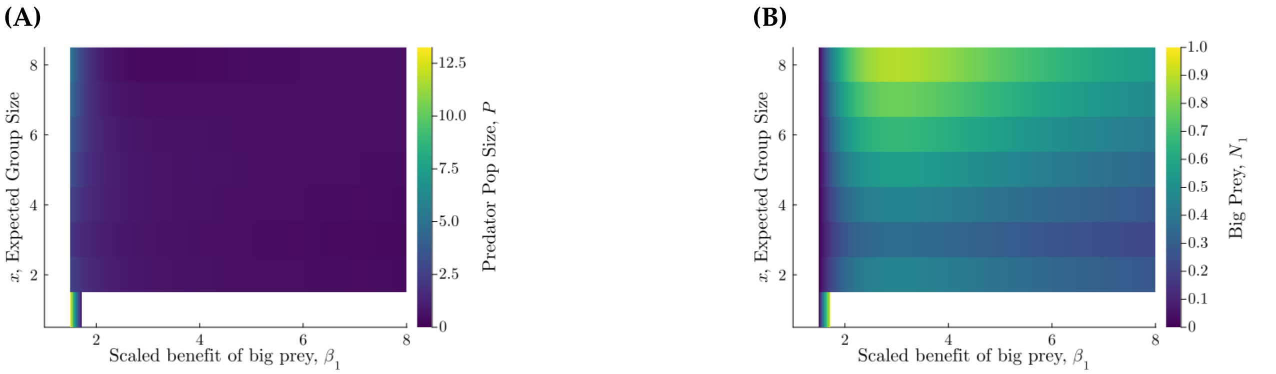


**Figure E.8:** The effect of group size and the benefit of big prey on the equilibrium **(A)** predator and **(B)** prey population sizes if only big prey are present. In panels **(A), (B)**, white space indicates that predators go extinct. The functional response is Type I, with $H_1(x) = 0$. Parameters follow allometric scaling, (see eq. 8), where $\beta_2 = 1$, $\alpha_1(1) = 0.95, \alpha_2 = 0.05, s_1 = 2, \eta_2 = 0.6$, and $\beta_2 = 1.0$ (note that we specify small prey-associated parameters in order to calculate big prey parameters using allometric scaling, but small prey is not present).

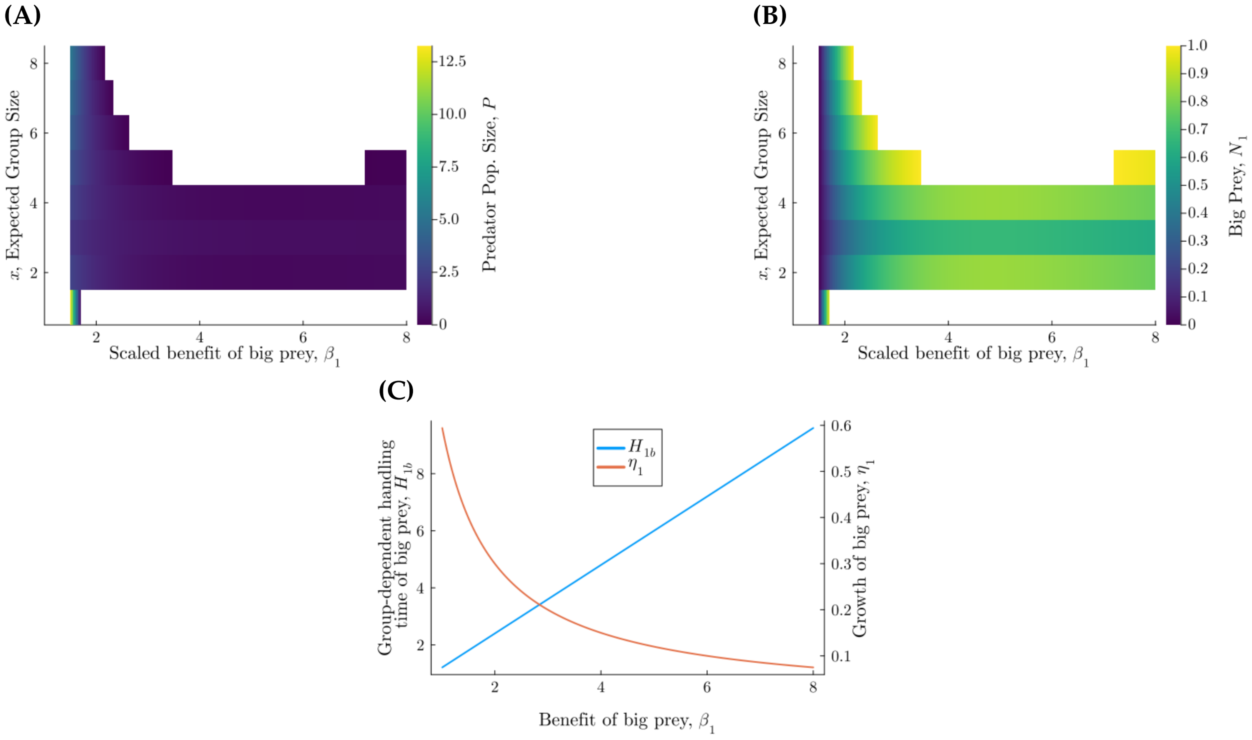


**Figure E.9:** The effect of group size and the benefit of big prey on the equilibrium **(A)** predator and **(B)** prey population sizes if only big prey are present. In panels **(A), (B)**, white space indicates that predators go extinct. Parameters follow allometric scaling, (see eq. 8), where $\beta_2 = 1$, $H_{2b} = 1.0$. $\alpha_1(1) = 0.95$, $\alpha_2 = 0.05$, $s_1 = 2$, $\eta_2 = 0.6$, and $\beta_2 = 1.0$ (note that we specify small prey-associated parameters in order to calculate big prey parameters using allometric scaling, but small prey is not present). The values of group-dependent handling time, $H_{1b}$ and scaled population growth of big prey, $\eta_1$, are in panel **(C)**.

##### E.2.2 *Two Prey Populations*

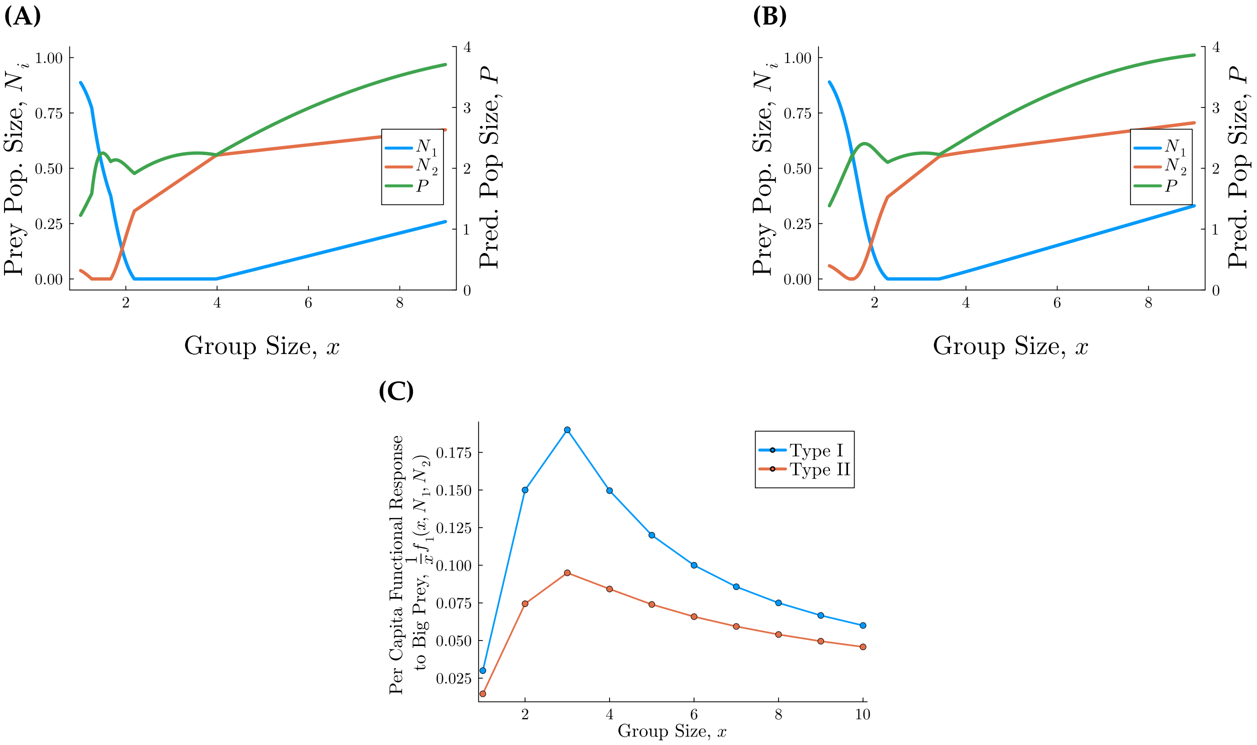


**Figure E.10:** The effect of group size on the equilibrium predator and prey population sizes, without group dynamics (predators are in groups of the same size). Curves show stable equilibrium values of predator population size, i.e., $P$ (green), and scaled big prey and small prey population sizes, i.e., $N_1, N_2$ (blue, red, respectively), for **A)** a Type I functional response, i.e., $H_1(x) = H_2(x) = 0$ and **B)** a Type II functional response, i.e., $H_1(x), H_2(x) > 0$. For both panels, the parameters use scaling laws (eq. 8), where the mass ratio of prey is $\beta_1/\beta_2 = 1.8$, for $\alpha_1(1) = 0.95, \alpha_2 = 0.05, s_1 = 2, \eta_2 = 0.6$, and $\beta_2 = 1.0$. For panel **B)**, $H_{1a} = H_{2a} = 0$ and $H_{2b} = 1.0$. Panel **C)** Shows the per-capita functional response for the parameters in Panels **A, B**, which are blue and red, respectively.

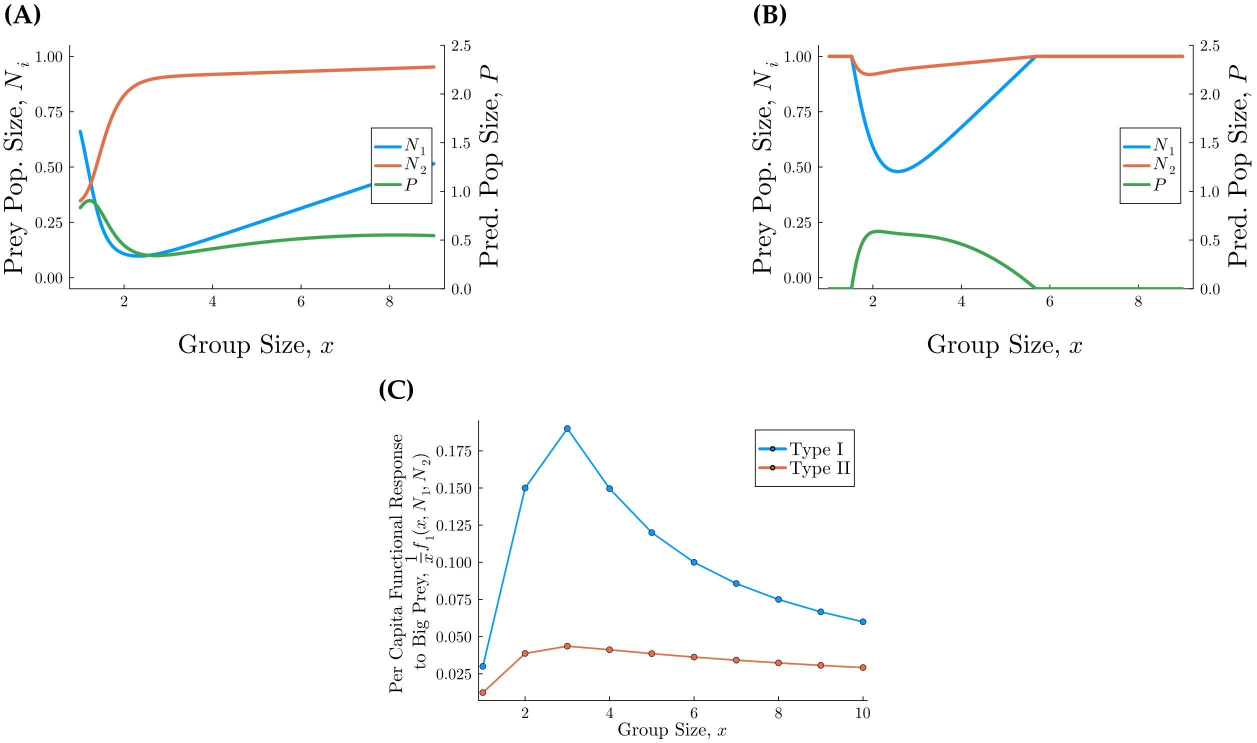


**Figure E.11:** The effect of group size on the equilibrium predator and prey population sizes, without group dynamics (predators are in groups of the same size). Curves show stable equilibrium values of predator population size, i.e., $P$ (green), and scaled big prey and small prey population sizes, i.e., $N_1, N_2$ (blue, red, respectively), for **A)** a type I functional response, i.e., $H_1(x) = H_2(x) = 0$ and **B)** a type II functional response, i.e., $H_1(x), H_2(x) > 0$. For both panels, the parameters use scaling laws (eq. 8), where the mass ratio of prey is $\beta_1/\beta_2 = 8$, for $\alpha_1(1) = 0.95, \alpha_2 = 0.05, s_1 = 2, \eta_2 = 0.6$, and $\beta_2 = 1.0$. For panel **B)**, $H_{1a} = H_{2a} = 0$ and $H_{2b} = 1.0$. Panel **C)** Shows the per-capita functional response for the parameters in Panels **A, B**, which are blue and red, respectively.

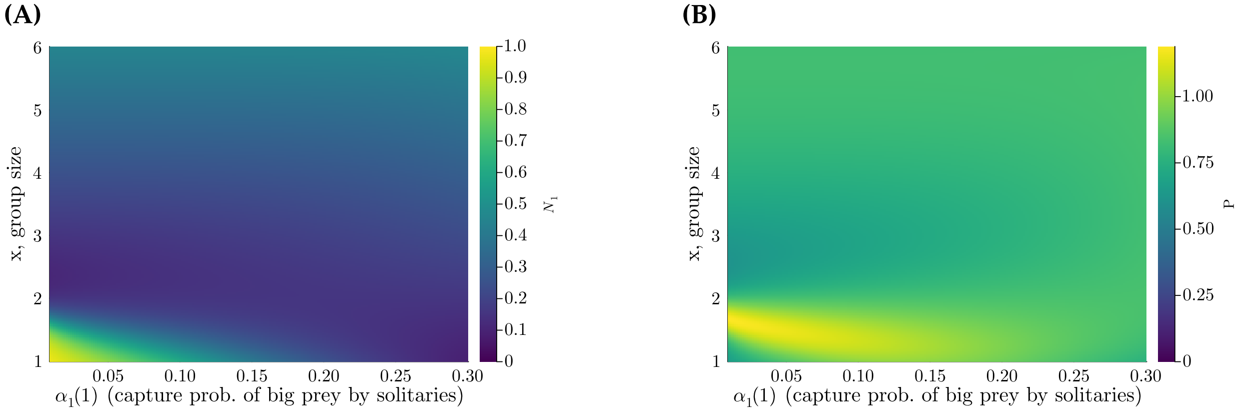


**Figure E.12:** Sensitivity analysis of the system with population dynamics but without group dynamics to the group size, $x$, (vertical axis), and the capture probability of big prey by solitaries, $\alpha_1(1)$, where the functional response is type I, i.e., the handling times are $H_1(x) = H_2(x) = 0$. The heatmaps show the stable coexistence equilibrium values of **A)** $N_1$, (scaled) big prey population size, and **B)** $P$, predator population size. The parameters are $s = 2$, $A_1 = 0.6, A_2 = 0.5, \beta_2 = 1.0, \eta_2 = 0.6, \alpha_2(1) = 0.95$, and the parameters $\beta_2, \eta_1$ are found using allometric scaling (see eq. 8) with the mass ratio being $\beta_1/\beta_2 = 4.0$. Although not shown, there is very little visible effect of $\alpha_1(1)$ on $N_2$, the small prey scaled population size.

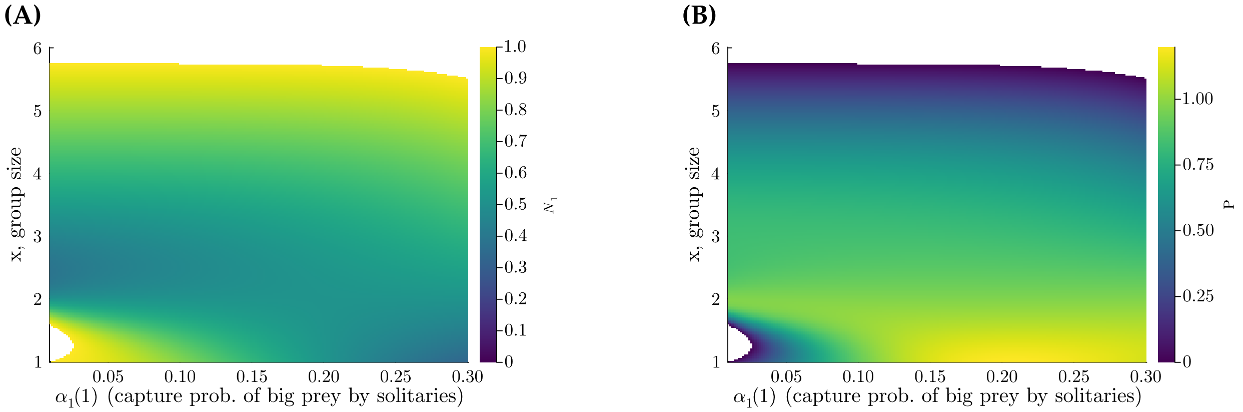


**Figure E.13:** Sensitivity analysis of the system with population dynamics but without group dynamics to the group size, $x$, (vertical axis), and the capture probability of big prey by solitaries, $\alpha_1(1)$, where the functional response is type II, i.e., the handling times $H_1(x), H_2(x) > 0$. The heatmaps show the stable coexistence equilibrium values of **A)** $N_1$, (scaled) big prey population size, and **B)** $P$, predator population size. The parameters are $s = 2$, $A_1 = 0.6, A_2 = 0.5, \beta_2 = 1.0, \eta_2 = 0.6, \alpha_2(1) = 0.95, H_{1a} = H_{2a} = 0$, and $H_{2b} = 1.0$. The the parameters $\beta_2, \eta_1, H_{1b}$ are found using allometric scaling (see eq. 8) with the mass ratio being $\beta_1/\beta_2 = 4.0$. Areas without a stable coexistence equilibrium are not colored in. Although not shown, there is very little visible effect of $\alpha_1(1)$ on $N_2$, the small prey scaled population size.

#### E.4 Apparent Competition, No Group Dynamics

Apparent competition occurs if $\frac{\partial N_2}{\partial N_1} < 0$. Say all predators are in groups of the same size, $x$, which is constant. For $\frac{dP}{dT}$ defined in eq. 12a, we can write

$$\frac{1}{P}\frac{\mathrm{d}P}{\mathrm{d}T} = -\tilde{\delta} + \frac{1}{x}\sum_{i=1,2} \beta_i \tilde{f}_i(x, N_1, N_2) = U_P(N_1, N_2)$$

Along the predator isocline, as in Holt (1977),

$$0 = \frac{\partial U_P}{\partial N_1} + \frac{\partial U_P}{\partial N_2}\frac{\partial N_2}{\partial N_1}$$

so

$$\frac{\partial N_2}{\partial N_1} = -\left(\frac{\partial U_P/\partial N_1}{\partial U_P/\partial N_2}\right) = -\left(\frac{\beta_1 \frac{\partial \tilde{f}_1}{\partial N_1} + \beta_2 \frac{\partial \tilde{f}_2}{\partial N_1}}{\beta_1 \frac{\partial \tilde{f}_1}{\partial N_2} + \beta_2 \frac{\partial \tilde{f}_2}{\partial N_2}}\right),$$

where the partial derivatives of the functional responses with respect to $N_1, N_2$ are in eqs. E.45. Substituting these into the equation above, we have

$$\frac{\partial N_2}{\partial N_1} = -\frac{\beta_1 A_1 \alpha_1(x)\left(1 + \alpha_2 H_2(x) N_2\right) - \beta_2 A_2 \alpha_1(x) \alpha_2 H_1(x) N_2}{-\beta_1 A_1 \alpha_1(x) \alpha_2 H_2(x) N_1 + \beta_2 A_2 \alpha_2 \left(1 + \alpha_1 H_1(x) N_1\right)},$$

which simplifies to,

$$\frac{\partial N_2}{\partial N_1} = -\left(\frac{\alpha_1(x)}{\alpha_2}\right)\left(\frac{A_1 \frac{\beta_1}{\beta_2} + \alpha_2 N_2 \left(\frac{\beta_1}{\beta_2} A_1 H_2(x) - A_2 H_1(x)\right)}{A_2 - \alpha_1(x) N_1 \left(A_1 \frac{\beta_1}{\beta_2} H_2(x) - A_2 H_1(x)\right)}.\right) \tag{E.48}$$

**Result 6.** *Big prey and small prey exert apparent competition on each other, i.e.,* $\frac{\partial N_2}{\partial N_1}, \frac{\partial N_1}{\partial N_2} < 0$, *if all predators are solitary and parameters follow the scaling relationships from Weitz and Levin (2006) based on eq. 8.*

*Proof.* If we assume the scaling assumptions in eq. 8, then eq. E.48 becomes

$$\frac{\partial N_2}{\partial N_1} = -\left(\frac{\alpha_1(1)}{\alpha_2}\right)\left(\frac{\beta_1}{\beta_2}\right)\left(\frac{A_1}{A_2}\right) < 0. \tag{E.49}$$

Note that apparent competition of small prey onto big prey is just the inverse. Thus there is apparent competition, and not facilitation, in both directions between big prey and small prey . ☐

**Result 7.** *For all predators in groups of size x and the populations at a stable coexistence equilibrium, then big prey and small prey exert apparent competition on each other if the following conditions are true: (1) parameters follow the scaling relationships from Weitz and Levin (2006) based on eq. 8; and (2)* $H_{1a} = H_{2a} = 0$, .

*Proof.* The proof is the same as the proof for Result 6. ☐

### F Population and Group Dynamics Together

#### *F.1 Equilibria*

The predator population, big prey, and small prey nullcines are obtained by setting the right sides of eqs. 12 equal to 0, i.e.,

$$0 = -\tilde{\delta} P + \sum_x g_x \sum_{i=1,2} \frac{A_i \beta_i \alpha_i(x) N_i}{1 + \alpha_1(x) H_1(x) N_1 + \alpha_2 H_2(x) N_2} \tag{F.50a}$$

$$0 = \eta_i (1 - N_i) - \sum_x \frac{A_i \alpha_i(x) g_x}{1 + \alpha_1(x) H_1(x) N_1 + \alpha_2 H_2(x) N_2} \text{ for } i = 1, 2, \tag{F.50b}$$

where eqs. F.50b are true for $i = 1, 2$ if $N_1 > 0$ or $N_2 > 0$, respectively. The null-clines for the time dynamics of $g_1, g_2, \cdots, g_{x_m}$ are obtained by setting the right sides of eqs. A.2a-d equal to 0, where $l = \phi = 1$ (so leaving and fusing of singletons have the same rate constants as joining of groups).

Note that because $\tilde{W}(x) = \frac{1}{x} \left( \beta_1 f_1(x, N_1, N_2) + \beta_2 f_2(x, N_1, N_2) \right)$ and $P = \sum_x x g_x$, from eq. F.50a,

$$0 = \sum_x x g_x \left[ \tilde{W}(x) - \tilde{\delta} \right] . \tag{F.51}$$

At the intersection of both prey non-zero nullclines and the predator population nullcline (eqs. F.50a and b),

$$\tilde{\delta}P = \sum_{i=1,2} \beta_i \eta_i N_i (1 - N_i). \tag{F.52}$$

The equilibrium must be solved numerically.

#### F.2 Local Stability

For $\vec{g} = (g_1, g_2, \cdots, g_{x_m})$, let $U_i(N_1, N_2, \vec{g})$ be the right sides of eq 12b,c for $i = 1, 2$ and $l = \phi = 1$. Let $\frac{\partial g_x}{\partial t} = Q_x(N_1, N_2, \vec{g})$ for $x = 1, 2, \cdots, x_m$, i.e., the right sides of eqs. A.2a-d, after replacing $\tau$ with $T_g$, $W(x)$ with $\tilde{W}(x)$, $\delta$ with $\tilde{\delta}$, and dividing by $T_g$.

Finding the partial derivatives to get the Jacobian:

$$\frac{\partial U_i}{\partial N_i} = \eta_i - 2\eta_i N_i - \sum_x g_x \frac{\partial \tilde{f}_i(x)}{\partial N_i}, \quad \frac{\partial U_i}{\partial N_j} = -\sum_x g_x \frac{\partial \tilde{f}_i(x)}{\partial N_j}, \quad \text{and } \frac{\partial U_i}{\partial g_x} = -\tilde{f}_i(x, N_1, N_2), \tag{F.53}$$

where $\frac{\partial t f_i}{\partial N_i}, \frac{\partial t f_i}{\partial N_j}$ are defined in eq. E.45. Also recall that the partial derivatives of fitness with respect to both prey types, $\frac{\partial \tilde{W}}{\partial N_i}$, are defined in eq. E.46.

To find the partial derivatives of $\frac{\partial g_x}{\partial t}$ with respect to $N_1, N_2$, we need the partial derivatives of the best response function. From the chain rule

$$\frac{\partial S(x,y)}{\partial N_i} = \frac{\partial S(x,y)}{\partial \tilde{W}(x)} \frac{\partial \tilde{W}(x)}{\partial N_i} + \frac{\partial S(x,y)}{\partial \tilde{W}(y)} \frac{\partial W(y)}{\partial N_i}$$

and because

$$\frac{\partial S(x,y)}{\partial W(x)} = dS(x,y)\left(1 - S(x,y)\right) \quad \text{and} \quad \frac{\partial S(x,y)}{\partial W(y)} = -dS(x,y)\left(1 - S(x,y)\right),$$

we have

$$\frac{\partial S(x,y)}{\partial N_i} = dS(x,y)\left(1 - S(x,y)\right)\left[\frac{\partial W(x)}{\partial N_i} - \frac{\partial W(y)}{\partial N_i}\right].$$

Then $\frac{\partial S(x,1)}{\partial N_i} = -\frac{\partial S(1,x)}{\partial N_i}$.

Then, for $\frac{\partial g_1}{\partial t}$ from eq. A.2a,

$$\frac{\partial Q_1}{\partial N_i} = \frac{1}{T_g}\left\{2g_2\frac{\partial S(1,2)}{\partial N_i} + \sum_{x=2}^{x_m}\left[xg_x\frac{\partial S(1,x)}{\partial N_i} - g_1 g_{x-1}\frac{\partial S(x,1)}{\partial N_i}\right]\right\} + x_m g_{x_m}\frac{\partial \tilde{W}(x_m)}{\partial N_i} - g_1\frac{\partial \tilde{W}(1)}{\partial N_i},$$

$$= \frac{1}{T_g}\left\{2g_2\frac{\partial S(1,2)}{\partial N_i} + \sum_{x=2}^{x_m}\frac{\partial S(1,x)}{\partial N_i}\left[xg_x + g_1 g_{x-1}\right]\right\} + x_m g_{x_m}\frac{\partial \tilde{W}(x_m)}{\partial N_i} - g_1\frac{\partial \tilde{W}(1)}{\partial N_i}, \tag{F.54}$$

and

$$\frac{\partial Q_1}{\partial g_1} = \frac{1}{T_g}\left\{-2g_1 S(2,1) - \sum_{x=3}^{x_m} g_{x-1}S(x,1)\right\} - \tilde{W}(1) - \tilde{\delta}, \tag{F.55}$$

$$\frac{\partial Q_1}{\partial g_2} = \frac{1}{T_g}\left[4S(1,2) - g_1 S(3,1)\right] + 2\tilde{\delta} \tag{F.56}$$

$$\frac{\partial Q_1}{\partial g_x} = \frac{1}{T_g}\left[xS(1,x) - g_1 S(x+1,1)\right] \quad \text{for } 2 < x < x_m \tag{F.57}$$

$$\frac{\partial Q_1}{\partial g_{x_m}} = \frac{1}{T_g}x_m S(1,x_m) + x_m\tilde{W}(x_m). \tag{F.58}$$

From eq. A.2b (after non-dimensionalizing and dividing by $T_g$),

$$\frac{\partial Q_2}{\partial N_i} = \frac{1}{T_g}\left\{-\frac{\partial S(1,2)}{\partial N_i}\left[2g_2 + \frac{1}{2}g_1^2\right] + \frac{\partial S(1,3)}{\partial N_i}\left[3g_3 + g_1 g_2\right]\right\}$$
$$+ g_1\frac{\partial \tilde{W}(1)}{\partial N_i} - 2g_2\frac{\partial \tilde{W}(2)}{\partial N_i}, \tag{F.59}$$

and

$$\frac{\partial Q_2}{\partial g_1} = \frac{1}{T_g}\left[g_1 S(2,1) - g_2 S(3,1)\right] + \tilde{W}(1) \tag{F.60}$$

$$\frac{\partial Q_2}{\partial g_2} = -\frac{1}{T_g}\left[2S(1,2) + g_1 S(3,1)\right] - 2\tilde{W}(2) - 2\tilde{\delta} \tag{F.61}$$

$$\frac{\partial Q_2}{\partial g_3} = \frac{3}{T_g}S(1,3) + 3\tilde{\delta} \tag{F.62}$$

$$\frac{\partial Q_2}{\partial g_x} = 0 \quad \text{for } x > 3. \tag{F.63}$$

From eq. A.2c, for $2 < x < x_m$,

$$\frac{\partial Q_x}{\partial N_i} = \frac{1}{T_g}\left\{\frac{\partial S(1,x+1)}{\partial N_i}\left[(x+1)g_{x+1}+g_1g_x\right] - \frac{\partial S(1,x)}{\partial N_i}\left[xg_x+g_1g_{x-1}\right]\right\} + (x-1)g_{x-1}\frac{\partial \tilde{W}(x-1)}{\partial N_i} - xg_x\frac{\partial \tilde{W}(x)}{\partial N_i}, \quad \text{(F.64)}$$

and

$$\frac{\partial Q_x}{\partial g_1} = \frac{1}{T_g}\left[g_{x-1}S(x,1) - g_xS(x+1,1)\right] \quad \text{(F.65)}$$

$$\frac{\partial Q_x}{\partial g_{x-1}} = \frac{1}{T_g}g_1S(x,1) + (x-1)\tilde{W}(x-1) \quad \text{(F.66)}$$

$$\frac{\partial Q_x}{\partial g_x} = -\frac{1}{T_g}\left[xS(1,x)+g_1S(x+1,1)\right] - x\tilde{W}(x) - x\tilde{\delta} \quad \text{(F.67)}$$

$$\frac{\partial Q_x}{\partial g_{x+1}} = \frac{1}{T_g}(x+1)S(1,x+1) + \tilde{\delta}(x+1) \quad \text{(F.68)}$$

$$\frac{\partial Q_x}{\partial g_y} = 0 \quad \text{for } y \neq 1, x-1, x, x+1. \quad \text{(F.69)}$$

Also from eq. A.2d, for $x = x_m$,

$$\frac{\partial Q_{x_m}}{\partial N_i} = -\frac{1}{T_g}\frac{\partial S(1,x_m)}{\partial N_i}\left[x_mg_{x_m}+g_1g_{x_m-1}\right] + (x_m-1)g_{x_m-1}\frac{\partial \tilde{W}(x_m-1)}{\partial N_i}, \quad \text{(F.70)}$$

and

$$\frac{\partial Q_{x_m}}{\partial g_1} = \frac{1}{T_g}g_{x_m-1}S(x_m,1) \quad \text{(F.71)}$$

$$\frac{\partial Q_{x_m}}{\partial g_{x_m-1}} = \frac{1}{T_g}g_1S(x_m,1) + (x_m-1)\tilde{W}(x_m-1) \quad \text{(F.72)}$$

$$\frac{\partial Q_{x_m}}{\partial g_{x_m}} = -\frac{1}{T_g}x_mS(1,x_m) - x_m\tilde{\delta} \quad \text{(F.73)}$$

$$\frac{\partial Q_{x_m}}{\partial g_y} = 0 \quad \text{for } y \neq 1, x_m-1, x_m. \quad \text{(F.74)}$$

#### F.3 *One Prey Population, Group and Population Dynamics Merged*

The Jacobian of this system is the Jacobian of the one-predator, two-prey system without the second column and row, with $N_2 = 0$.

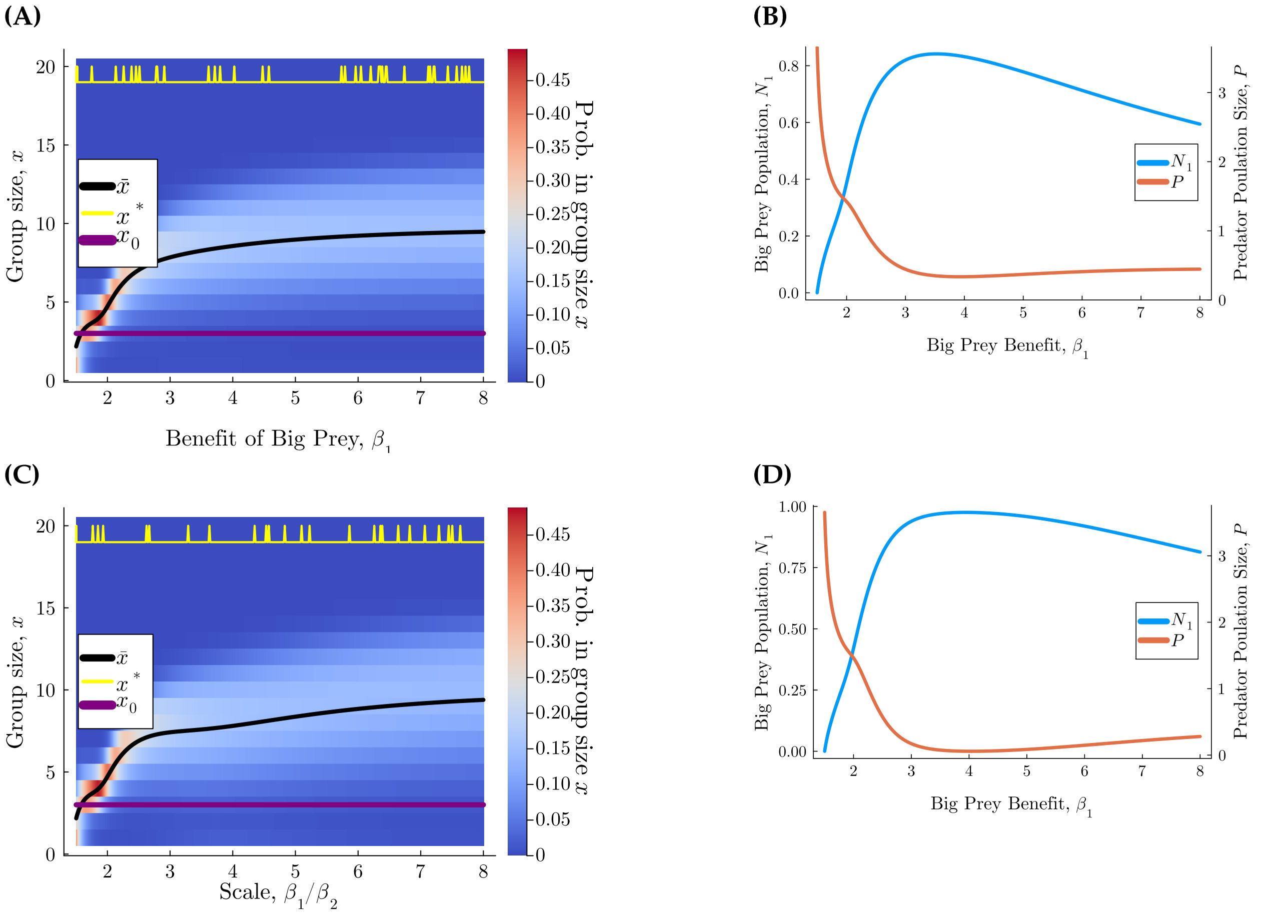


**Figure F.14:** Increasing the benefit of big prey increases group size while decreasing predator population size, for the system with group dynamics and only one prey type. The top row shows equilibrium values of **(A)** the probability of being in a group of size $x$ and **(B)** predator and big prey population sizes ($P$ and $N_1$, respectively), for a type I functional response, i.e., $H_1(x) = 0$. The bottom row shows equilibrium values of **(C)** the probability of being in a group of size $x$ and **(B)** $P$ and $N_1$, for a type II functional response, with $H_{2b} = 0.4$. Parameters are allometrically scaled, where $x_m = 20$, $\eta_2 = 0.6, \beta_2 = 1, A_1 = 0.6, A_2 = 0.5, T_x = 0.01, d = 100, \alpha_1(1) = 0.05, s_1 = 2$. In panels **(A)** and **(C)**, the purple line shows the group size that maximizes fecundity, the yellow line is the equilibrium group size predicted by Clark and Mangel (1986), and the black curve is the mean experienced group size. Spikes in the yellow line are artifacts of numerical instability because there are very small differences between $W(19)$, $W(20)$, and $W(1)$.

#### F.4 Bifurcation Diagrams, Two Prey Populations with Group Dynamics

In the following bifurcation diagrams, we use numerical continuation, and plot each subsequent branch found through numerical continuation with a different color. Thus, in one figure, one color indicates one branch (but different variables are shown in different panels due to the high dimensionality of the system) (Figs. F.15, F.16, F.17).

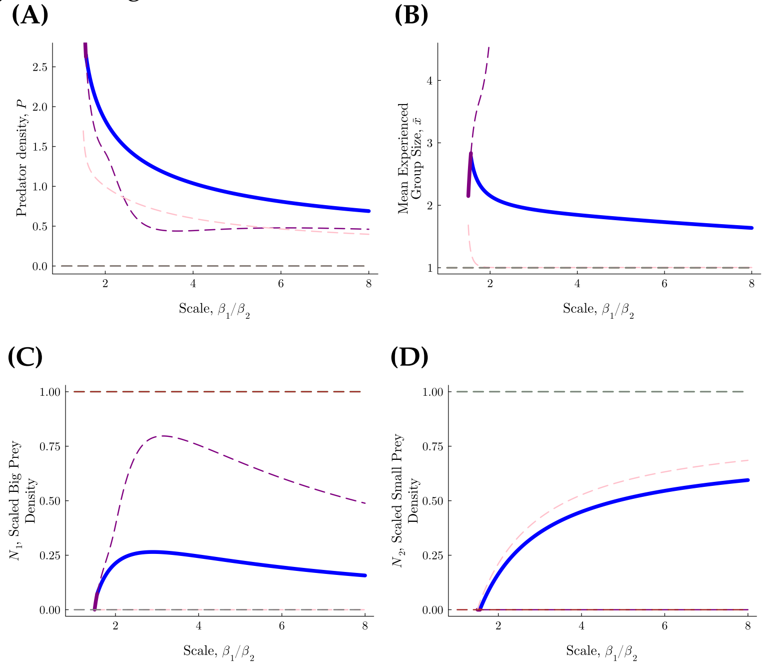


**Figure F.15:** Bifurcation diagrams of the system in relation to $\beta_1/\beta_2$ for a type 1 functional response $H_1(x) = H_2(x) = 0$) in which prey growth rate and benefit are scaled by bodymass such that $\beta_1/\beta_2 = \eta_1/\eta_2$. The maximum group size is $x_m = 5$, where the parameters are $\eta_2 = 0.6, \beta_2 = 1, A_1 = 0.6, A_2 = 0.5, T_x = 0.01, d = 100, \alpha_1(1) = 0.05, s_1 = 2$, and $\alpha_2(x) = 0.95$ is constant.. Stable equilibria branches are marked with solid, thick lines, and unstable branches are marked by dashed lines. The branches were found through numerical continuation, and each branch is colored a separate color. Note that mean experienced group size, $\bar{x} = 1$ if predators are extinct.

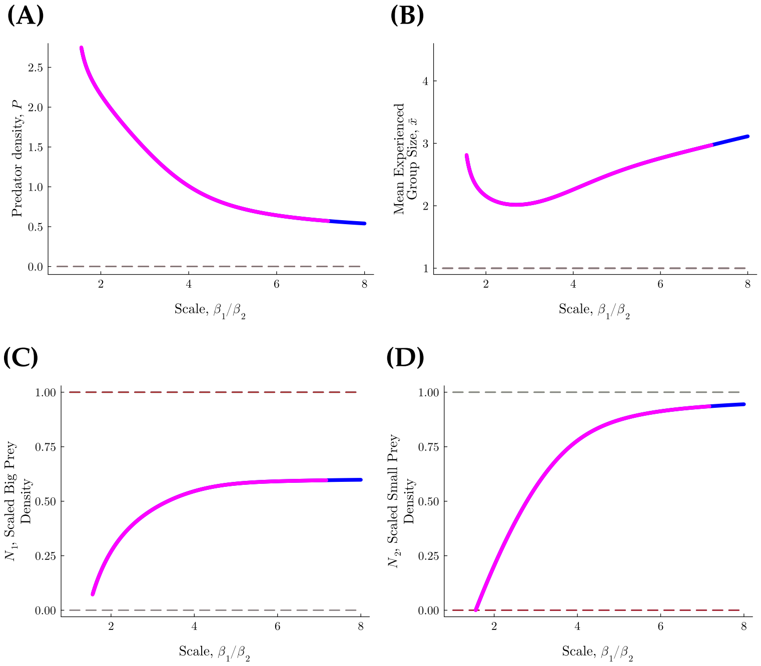


**Figure F.16:** Bifurcation diagrams of the system in relation to $\beta_1/\beta_2$, for the system in which $H_1, H_2, \beta_1, \beta_2, \eta_1, \eta_2$ are scaled by prey body mass such that $\beta_1/\beta_2 = \eta_1/\eta_2 = A_2H_1/(A_1H_2)$. The maximum group size is $x_m = 5$, and the parameters are $\eta_2 = 0.6, H_{1a} = H_{2a} = 0, H_{2b} = 1, \beta_2 = 1, A_1 = 0.6, A_2 = 0.5, d = 100, T_g = 0.01, \alpha_1(1) = 0.05, s_1 = 2$, and $\alpha_2(x) = 0.95$ is constant.. Stable equilibria branches are marked with solid, thick lines, and unstable branches are marked by dashed lines. The branches were found through numerical continuation, and each branch is colored a separate color. Note that mean experienced group size, $\bar{x} = 1$ if predators are extinct.

##### F.4.1 *Group Dynamics Much Faster than Population Dynamics, Two Prey Present*

Here we investigate why numerical continuation shows bistability of a coexistence equilibrium and a predator extinction equilibrium if group formation is very fast (Fig. F.17).

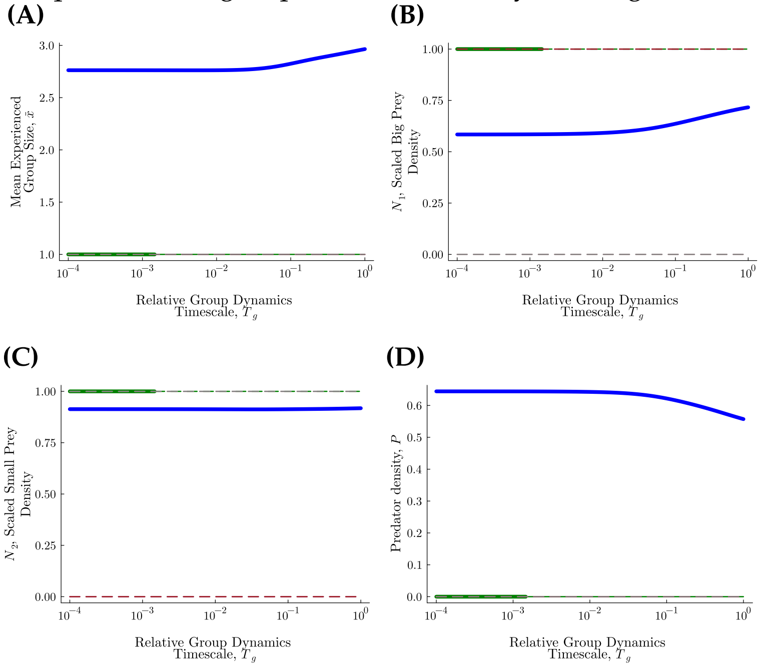


**Figure F.17:** Bifurcation diagrams of the system in relation to $T_g$, shown on a log scale, for the system in which $H_1, H_2, \beta_1, \beta_2, \eta_1, \eta_2$ are scaled by prey body mass such that $\beta_1/\beta_2 = \eta_1/\eta_2 = A_2H_1/(A_1H_2) = 6$. The maximum group size is $x_m = 5$, and the parameters are $\eta_2 = 0.6, H_{1a} = H_{2a} = 0, H_{2b} = 1, \beta_2 = 1, A_1 = 0.6, A_2 = 0.5, d = 100, T_x = 0.01, \alpha_1(1) = 0.05, s_1 = 2$, and $\alpha_2(x) = 0.95$ is constant. . Stable equilibria branches are marked with solid, thick lines, and unstable branches are marked by dashed lines. The branches were found through numerical continuation, and each branch is colored a separate color. Note that mean experienced group size, $\bar{x} = 1$ if predators are extinct.

If predators are extinct, substituting $\vec{g} = 0$ into the Jacobian with components defined in eqs. F.54 - F.74, the eigenvalues are $-\eta_1, -\eta_2$ and the solutions of the characteristic polynomial of an $x_m \times x_m$ jacobian, $M$ with components defined by eqs. F.55 - F.74, i.e., the partial derivatives of $Q_i(N_1, N_2, \vec{g})$ with respect to $\vec{g}$. This matrix, $M$, is zero everywhere except the top row, the lower diagonal, the diagonal, and the upper diagonal. Let $J = T_gM$ and let $\pi_x = xS(1,x)$. . The top row of $J$ is

$$\left(T_g(-\tilde{W}(1) + \tilde{\delta}), 2\pi_2 + 2T_g\tilde{\delta}, \pi_3, \pi_4, \cdots, \pi_{x_m-1}, \pi_{x_m} + x_mT_g\tilde{W}(x_m)\right)^T,$$

the lower diagonal is

$$T_g(\tilde{W}(1), 2\tilde{W}(2), \cdots, (x_m - 1)\tilde{W}(x_m - 1))^T,$$

the diagonal is

$$Diag(J) = \Big( T_g(-\tilde{W}(1) + \tilde{\delta}), -\pi_2 - 2T_g\left(\tilde{W}(2) + \tilde{\delta}\right), -\pi_3 - 3T_g\left(\tilde{W}(3) + \tilde{\delta}\right), \\ \cdots, -\pi_{x_m - 1} - (x_m - 1)T_g\left(\tilde{W}(x_m - 1) + \tilde{\delta}\right), -\pi_{x_m} - x_m T_g \Big)^T, \quad \text{(F.75)}$$

and the upper diagonal is

$$\left(2\pi_2 + 2T_g\tilde{\delta}, \pi_3 + 3T_g\tilde{\delta}, \pi_4 + 4T_g\tilde{\delta}, \cdots \pi_{x_m} + x_m T_g\tilde{\delta}\right).$$

For $T_g$ small enough, the lower diagonal can be approximated as the zero vector. Then this jacobian is upper triangular, and the eigenvalues are the diagonal elements listed in eq. F.75. Thus for $T_g$ small enough, the equilibrium in which $P = 0$ and $N_1 = N_2 = 1$ is stable if $\tilde{W}(1) > \tilde{\delta}$.

#### F.5 Sensitivity Analysis to $\alpha_1(1)$

As shown in Figs. F.18, F.19, for both a type I and type II functional response, making $\alpha_1(1)$ very small increases the mean experienced group size at equilibrium, $\bar{x}$, especially if the functional response is type II. Furthermore, there is a non-monotonic relationship between $\alpha_1(1)$ and prey population sizes, with big prey and small prey maximized and minimized, respectively, at $\alpha_1$ around 0.17, although overall the effect of capture probability by solitaries on the prey populations is not large.

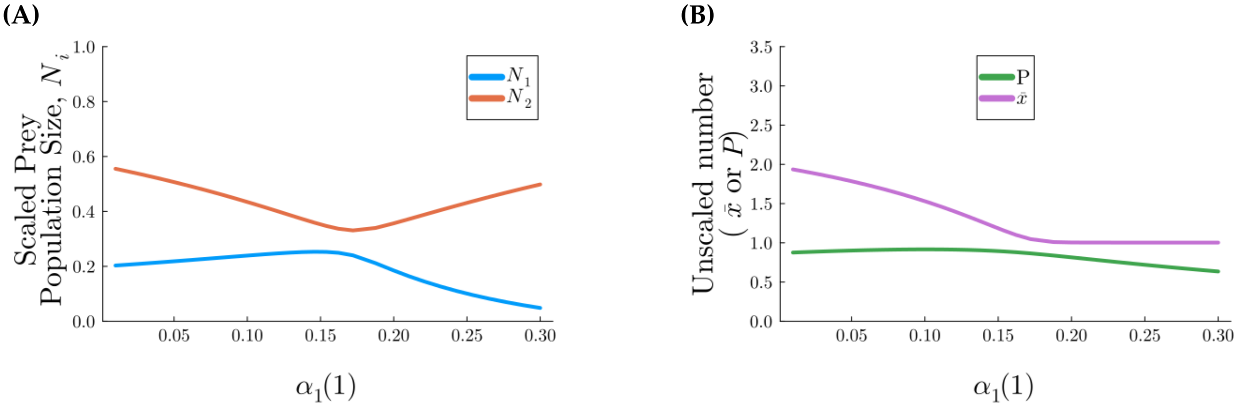


**Figure F.18:** Sensitivity analysis of the system with population dynamics and group dynamics to the capture probability of big prey by solitaries, $\alpha_1(1)$, where the functional response is type I, i.e., the handling times $H_1(x), H_2(x) = 0$. The curves show the stable coexistence equilibrium values. Panel textbfA) shows big prey and small prey (scaled) population sizes, i.e, $N_1$ and $N_2$, respectively. Panel **B)** focuses on predators, showing $P$, the predator population size, and $\bar{x}$, the mean experienced group size. The parameters are $s_1 = 2$, $A_1 = 0.6, A_2 = 0.5, \beta_2 = 1.0, \eta_2 = 0.6$, and $\alpha_2(1) = 0.95$. The the parameters $\beta_2$ and $\eta_1$ are found using allometric scaling (see eq. 8) with the mass ratio being $\beta_1/\beta_2 = 5.0$.

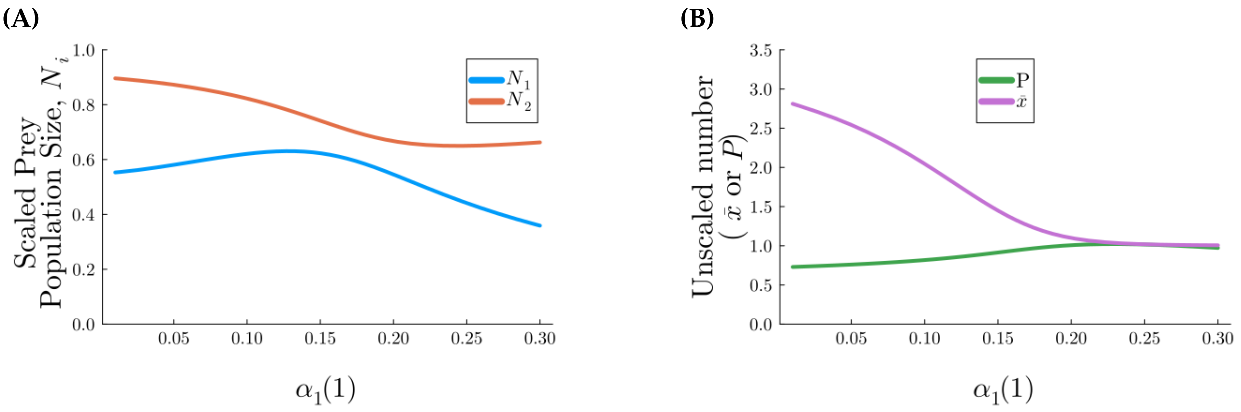


**Figure F.19:** Sensitivity analysis of the system with population dynamics and group dynamics to the capture probability of big prey by solitaries, $\alpha_1(1)$, where the functional response is type I, i.e., the handling times $H_1(x), H_2(x) > 0$. The curves show the stable coexistence equilibrium values. Panel textbfA) shows big prey and small prey (scaled) population sizes, i.e, $N_1$ and $N_2$, respectively. Panel **B)** focuses on predators, showing $P$, the predator population size, and $\bar{x}$, the mean experienced group size. The parameters are $s_1 = 2, A_1 = 0.6, A_2 = 0.5, \beta_2 = 1.0, \eta_2 = 0.6, H_{1a} = H_{2a} = 0, H_{2b} = 1.0$, and $\alpha_2(1) = 0.95$. The the parameters $\beta_2, H_{1b}$ and $\eta_1$ are found using allometric scaling (see eq. 8) with the mass ratio being $\beta_1/\beta_2 = 5.0$.

#### F.6 Apparent Competition

Recall that $Q_i(N_1, N_2, g_1, g_2, \cdots, g_{x_m}) = \frac{\partial g_i}{\partial T}$ and let $U_i(N_1, N_2, g_1, g_2, \cdots, g_{x_m}) = \frac{\mathrm{d}N_i}{\mathrm{d}T}$. Along the prey and group size distribution nullclines,

$$0 = \frac{\partial U_1}{\partial N_1} + \frac{\partial U_1}{\partial N_2}\frac{\partial N_2}{\partial N_1} + \sum_{x=1}^{x_m} \frac{\partial U_1}{\partial g_x}\frac{\partial g_x}{\partial N_1}$$

$$0 = \frac{\partial U_2}{\partial N_1} + \frac{\partial U_2}{\partial N_2}\frac{\partial N_2}{\partial N_1} + \sum_{x=1}^{x_m} \frac{\partial U_2}{\partial g_x}\frac{\partial g_x}{\partial N_1}$$

$$0 = \frac{\partial Q_i}{\partial N_1} + \frac{\partial Q_i}{\partial N_2}\frac{\partial N_2}{\partial N_1} + \sum_{x=1}^{x_m} \frac{\partial Q_i}{\partial g_x}\frac{\partial g_x}{\partial N_1} \quad \text{for } i = 1, 2, \cdots, x_m,$$

and we can solve this linear set of equations for $\frac{\partial N_2}{\partial N_1}$ by solving it for $\frac{\partial N_2}{\partial N_1}, \frac{\partial g_1}{\partial N_1}, \frac{\partial g_2}{\partial N_1}, \cdots, \frac{\partial g_{x_m}}{\partial N_1}$ numerically using matrix methods (The vector $(\frac{\partial N_2}{\partial N_1}, \frac{\partial g_1}{\partial N_1}, \frac{\partial g_2}{\partial N_1}, \cdots, \frac{\partial g x_m)}{\partial N_1})^T$ is the inverse of the Jacobian $M$ derived in Appendix Section F.2 without the first row and the first column, which we call $M_{-1}$, multiplied by the vector $-(\frac{\partial U_2}{\partial N_1}, \frac{\partial Q_1}{\partial N_1}, \frac{\partial Q_2}{\partial N_1}, \cdots, \frac{\partial Q_{x_m}}{\partial N_1})^T$).

$$
-\begin{pmatrix} \frac{\partial U_2}{\partial N_1} \\ \frac{\partial Q_1}{\partial N_1} \\ \frac{\partial Q_2}{\partial N_1} \\ \vdots \\ \frac{\partial Q_{x_m}}{\partial N_1} \end{pmatrix} = M_{-1} \begin{pmatrix} \frac{\partial N_2}{\partial N_1} \\ \frac{\partial g_1}{\partial N_1} \\ \frac{\partial g_2}{\partial N_1} \\ \vdots \\ \frac{\partial g_{x_m}}{\partial N_1} \end{pmatrix} . \tag{F.76}
$$

Symmetrically, to find the apparent competition of prey 2 on prey 1,

$$
-\begin{pmatrix} \frac{\partial U_1}{\partial N_2} \\ \frac{\partial Q_1}{\partial N_2} \\ \frac{\partial Q_2}{\partial N_2} \\ \vdots \\ \frac{\partial Q_{x_m}}{\partial N_2} \end{pmatrix} = M_{-2} \begin{pmatrix} \frac{\partial N_1}{\partial N_2} \\ \frac{\partial g_1}{\partial N_2} \\ \frac{\partial g_2}{\partial N_2} \\ \vdots \\ \frac{\partial g_{x_m}}{\partial N_2} \end{pmatrix} , \tag{F.77}
$$

where $M_{-2}$ is the jacobian without the 2nd row and 2nd column.
